# Variation in gut microbiome diversity and structure across host lifestyles in Thailand

**DOI:** 10.64898/2026.01.12.699019

**Authors:** Natchapon Srinak, Ratha-korn Vilaichone, Lucas Moitinho-Silva, Christopher H. Corzett, Christoph Kaleta, Eric Alm, Mathieu Groussin, Varocha Mahachai, Mathilde Poyet, Jan Taubenheim

## Abstract

Global comparisons have revealed marked shifts in gut microbiome diversity and composition with industrialization and urbanization. Whether such changes also arise at finer geographical scales, among neighboring populations adopting different lifestyles, remains largely unexplored. In this study, we retrieved 16S rRNA sequencing data and characterized the gut microbiomes of Thai populations with distinct lifestyles: urban residents from Bangkok, farmers from Tak province, and foragers from Phatthalung province. Our results reveal that microbiomes of all populations are primarily dominated by the families Prevotellaceae, Bacteroidaceae, and Lachnospiraceae, yet their specific abundances differ between populations. Diversity measurements, particularly Faith’s phylogenetic diversity, showed a descending trend from the rural to urban populations. Despite these differences, results suggested that most highly prevalent microbial genera were shared among all groups, suggesting the existence of a consistent shared microbiome within the Thai population, despite differing lifestyles. Further, association analysis shows that overall population-wide lifestyle was significantly associated with microbial community structure. Linking dietary habits and other host factors to taxa abundance using LASSO and linear modeling revealed population-specific microbiome-lifestyle associations, indicating that baseline microbial community composition determines microbiome variability in response to environmental changes. Overall, our study expands the scope of host–lifestyle–microbiome research and identifies associations between lifestyle and microbial features, underscoring the influence of lifestyle factors and baseline microbiomes on microbial compositional adaptation.

## Introduction

The human gut microbiome harbors approximately 100 times more genes (∼3-5 million) than the human genome itself (∼20,000 genes) (1–3). This immense genetic reservoir provides a wide range of functions that are closely associated with human health and fitness, involved in various key processes such as development, aging, and behavior (4–6). Although microbial functions tend to be diverse and relatively stable (7), industrialized lifestyle factors, such as diet (8,9), urban housing (10), hygiene practices (11), antibiotic exposure (12), or cesarean births (13), have significant effect on microbiome composition, affecting diversity, transmission, and persistence of certain microbial taxa (14). In contemporary microbiome studies, these conditions and microbiomes are treated as a ‘default’, even though disease prevalence and microbiome composition are both influenced by the degree of industrialization (15–17). However, our understanding of how industrialization affects microbes is limited, mostly because the majority of human populations studied live under industrialized conditions.

To understand how these industrialization-associated shifts arose, studies have characterized human gut microbiomes across major lifestyles from pre-industrialization to present time (18,19). The first major shift occurred with the transition from hunting and gathering to farming and herding, followed by industrialization and urbanization; each of these stages directly shaped the composition and diversity of gut microbial communities. Ancient hunter-gatherer communities, typically nomadic or semi-nomadic and organized in small groups, relied primarily on locally available resources without or with minimal manual processing, including wild plants and animals. Analyses of coprolites (fossilized feces) indicate that their microbiomes differ substantially from those of extant urban populations, yet they approximate those of present-day rural human communities (20). Studies of non-industrialized indigenous communities in Tanzania, Venezuela, and Papua New Guinea have shown that the genus *Prevotella* is enriched in their gut microbiomes, and that overall α-diversity is higher compared with industrialized populations (21–23). Adaptation of subsistence strategies from foraging to farming and herding shifted diet choices from diverse wild plants and animals to limited crops and domestic livestock. Microbiome studies in non-industrialized farmers and fishermen in rural Cameroon and Malaysia reported that this lifestyle has shown only a slight decrease in microbial diversity in comparison to foraging populations (24,25). However, it showed some changes in microbial composition; for example the increased abundance of *Bifidobacterium*, *Ruminococcus* and *Prevotella*. With increased industrialization, life became isolated from natural environments, and diets tend to be dominated by high amounts of simple carbohydrates and fats in (ultra-) processed foods accompanied by a reduction in fiber (17,26). In addition, industrialized lifestyles encompass different hygiene practices like frequent use of sanitizers and antibiotics, which have reshaped the urban microbial landscape (11,18). Consistently, studies on gut microbial diversity reported that the urban populations exhibit substantially lower diversity compared with populations depending on rural subsistence strategies (21,22,24,25,27–29) and that inter-population microbial distance was higher in industrialized populations (30).

These studies provide fundamental insights into how lifestyle shapes the human gut microbiome. However, most microbiome research focuses on urban caucasian individuals from industrialized countries and centers around medical applications, leaving many human populations underrepresented in microbiome science. Microbiomes from these under-sampled populations often differ substantially from those of industrialized cohorts (31), which limits the generalizability of existing findings (32). Moreover, studies examining how individual lifestyle factors shape microbial diversity or the abundance of specific taxa remain limited and can be context dependent. Characterizing microbiomes across diverse lifestyles in underrepresented populations is therefore essential to fill knowledge gaps, expand the global microbiome catalog, and establish baseline profiles that link lifestyle with health and disease.

In this study, we used 16S rRNA sequencing to characterize gut microbiomes from three Thai cohorts representing three distinct lifestyles: foragers from the Phatthalung province (self identified as “Maniq” people), farmers from the Tak province (self identified as “Hmong” people), and urban residents from Bangkok (self identified as “Thai” people). These samples are part of Global Microbiome Conservancy (GMbC; https://microbiomeconservancy.org/). The participants recruited from the Phatthalung province, are part of one of the few remaining human populations in the world that primarily subsists through foraging activities. While they have some access to products provided by health organizations, a previous study reported that they primarily rely on wild plants (such as yams) and animal resources (33,34). They also maintain traditional cultural practices, including the use of medicinal herbs, and the continuation of historic birth rituals (33). A recent genetic study indicated that they have admixed ancestry, showing East Asian-related genetic similarity to other Thai ethnic groups (35). The participants recruited from Tak province rely primarily on farming as their main subsistence strategy, including crop cultivation and livestock rearing (36). Historically, the Hmong migrated to Thailand approximately 200 years ago from northern Myanmar, northern Laos, and southern China (37). Prior to migration, they practiced agriculture, a subsistence pattern they have continued to maintain in Thailand. For the urban group, samples were collected from residents of Bangkok, a metropolitan region that has high population density and socioeconomic diversity (38). According to genome-wide analyses (39), most Thai populations (including urban residents of Bangkok) originate primarily from southern China migrants, with additional Austroasiatic and South Asian genetic admixture. Today, Bangkok represents a highly industrialized and urbanized city where lifestyles commonly involve the consumption of high-fat and high-sugar (ultra-) processed foods (40).

Here, we established baseline microbiome profiles for each lifestyle, identified uniquely enriched microbial taxa potentially linked to lifestyle. We further conducted association analyses between both composite lifestyle features and individual lifestyle variables with microbial features. We detected differences in microbial composition among the three lifestyles. However, 58% of genera were shared across all populations, and microbial prevalence was highly correlated between populations, suggesting that many high-prevalence genera may be common in the Thai microbiome regardless of host factors. Diversity analyses revealed the highest richness and phylogenetic diversity in samples from Phatthalung individuals, intermediate levels in Tak individuals, and the lowest in Bangkok microbiomes, which were instead characterized by high evenness and reduced phylogenetic breadth. β-diversity analyses confirmed significant separation between populations, with urban samples showing the greatest inter-individual variability. Differential abundance analysis identified 34 genera and 65 amplicon sequence variants (ASVs) distinguishing the microbiomes from the three populations. Lifestyle and dietary factors correlated with overall microbial structure and fine-scale analyses revealed population-specific associations between microbial abundances and food items. Although the Thai populations shared many microbial genera, their gut microbial communities differed. Urbanization and lifestyle diversification, and dietary differences, were associated with reduced microbial richness and phylogenetic diversity.

## Material and Methods

### Data collection

#### Participants, criteria and lifestyle collection

Samples were obtained from three geographically distinct cohorts, each representing a different lifestyle. The cohorts included: 47 urban, industrialized individuals recruited from Bangkok (mean age ± SD: 30.8 ± 6.22 years); 29 individuals from foraging communities located in Phatthalung Province (mean age ± SD: 37.3 ± 9.78 years); and 30 farmers residing in Tak Province (mean age ± SD: 30.9 ± 9.01 years). Participants were recruited as part of the cross-sectional Global Microbiome Conservancy cohort, which was designed to capture variation in geography, urbanization, subsistence and lifestyle. Eligible participants were adults who reported no symptoms of infectious or chronic disease at enrolment. Written informed consent was obtained in an appropriate local language. The study was approved in Thailand by the Human Research Ethics Committee of the Faculty of Medicine, Thammasat University No. 1 (Certificate of Approval 143/2019). Demographic and host data, including age, sex and anthropometric characteristics, were recorded together with information on lifestyle, environmental exposures and medication use. Lifestyle variables included household size, water source, contact with animals, tobacco use, physical activity, access to electricity, floor type, recent international travel and other early-life or customary factors, including mode of delivery, breastfeeding, purging practices and dietary data. Recent antibiotic and antiparasitic use was also documented. Habitual dietary intake was assessed using population-adapted food-frequency questionnaires in a semi-quantitative manner (ranging from 5 - very often to 1 - never). A summary of all host features is in Supplementary Table 1, while a full meta-data table is available in the recent GMbC publication (41). The interpretation of each variable score is provided in Supplementary Table 2. A standardized core questionnaire was supplemented with locally relevant foods, and the questionnaires were subsequently harmonized across populations to retain local dietary specificity while facilitating comparisons between study sites.

### Stool sample collection

Stool samples were collected between November and December 2019. Participants were provided with empty sterile 15 mL vials (Thermo Scientific™ Empty Vial) and instructed to deposit fecal material directly into the container. All samples were processed within one hour of defecation. Approximately 1 g of stool was transferred into a sterile vial containing 10 mL of RNAlater preservative. The preserved samples were shipped to the Massachusetts Institute of Technology (MIT, USA) and stored at −80 °C within the Global Microbiome Conservancy biobank for downstream analyses.

#### 16S rRNA extraction and sequencing

Bacterial genomic DNA was extracted from each fecal sample using the PowerSoil DNA Isolation Kit (Qiagen) according to the manufacturer’s instructions. The bacterial community composition was determined by sequencing the V3–V4 region of the 16S ribosomal RNA (rRNA) gene using an Illumina MiSeq platform at the Broad Institute of MIT and Harvard, following previously published protocols (42).

#### 16S rRNA identification

Paired-end 16S rRNA reads were first quality-checked using FastQC v0.12.1 (43) and MultiQC v1.25.1 (44). On average, the reads had a GC content of 51.92% and samples contained 46,934 reads (Supplementary Table 3), with most samples (197 out of 212) showing high Phred scores. Reads were then preprocessed using the filterAndTrim function in the DADA2 package (45), which removed low-quality reads with Phred scores below 11 (truncQ = 11) and filtered out reads with more than two expected errors (maxEE = 2). The cleaned reads were subsequently processed for taxonomic clustering and assignment using DADA2. Reads were clustered into amplicon sequence variants (ASVs), distinguishing sequences differing by as little as a single nucleotide. Taxonomic assignment was performed using a Naive Bayes classifier trained on the SILVA nr99 database (v138.1_wSpecies) (46). Taxonomic annotation and count can be found in Supplementary Table 4. Quality-checking steps were performed in Bash, while all other analyses were conducted in R v4.4.1. For more details see the extended method section.

### Statistical analysis

#### α-diversity and phylogenetic diversity

Shannon and Simpson diversity indices were computed using the vegan package v2.6.8 (47). Species richness for each sample was counted as the total number of unique ASVs with non-zero abundance. Later, species evenness, or Shannon equitability index was calculated as: Evenness = Shannon/ln(Richness).

For phylogenetic diversity, we calculated Faith’s Phylogenetic Diversity (Faith’s PD) and Phylogenetic Species Variability (PSV). To derive a phylogenetic tree, all ASV sequences were first extracted and converted to a FASTA file using Biostrings v2.74.1. Multiple sequence alignment (MSA) was performed using MUSCLE v5.1 (48), and the alignment was refined to retain highly conserved nucleotide blocks using Gblocks v0.91b (49). The phylogenetic tree was then reconstructed with FastTree v2.2 (50), and its midpoint was estimated using phangorn v2.12.1 (51). Faith’s PD and PSV were subsequently calculated with Picante v1.8.2 (52). The MSA, alignment refinement, and tree construction steps were performed in Bash, while all other analyses were conducted in R v4.4.1.

To test differences of diversities between populations (Bangkok, Tak, and Phatthalung), pairwise Wilcoxon tests were conducted using rstatix v0.7.2 (53). For more details see the extended method section.

#### β-diversity

β-diversity was assessed using Bray-Curtis dissimilarity, Aitchison distance, unweighted UniFrac, and weighted UniFrac. Bray-Curtis dissimilarity was calculated directly from the relative abundance matrix using the vegan package v2.6.8 (47), while unweighted and weighted UniFrac distances were computed in phyloseq v1.50.0 (54) using the same phylogenetic tree as in the phylogenetic diversity. For these three distance matrices, principal coordinates analysis (PCoA) was performed using ape v5.8 (55). Aitchison distance was computed by first transforming the relative abundance matrix using the centered log-ratio (CLR) transformation, followed by calculation of Euclidean distance with the compositions v2.0.8 (56). Principal component analysis (PCA) on the CLR-transformed matrix was performed using stats v4.4.1.

Multivariate dispersions, representing intra-population microbial composition variance, were calculated for the Bangkok, Tak, and Phatthalung samples using each β-diversity matrix with the vegan. Dispersions for each index were statistically compared using ANOVA implemented in the compositions. Additionally, pairwise PERMANOVA tests were performed for each population pair across all β-diversity indices using the pairwiseAdonis v0.4.1. P-values were adjusted using the Bonferroni method (p.adjust.m = “bonferroni”), and the number of permutations was set to 10,000 (perm = 10000). For more details see the extended method section.

#### Phylogenetic tree

For better visualization and interpretation, a phylogenetic tree at genus level was reconstructed. Firstly, representative ASVs for each genus were selected by highest relative abundance across samples. The process to reconstruct the tree was the same as described earlier.

#### Prevalence analysis

Prevalence of each microbial genus was calculated for each location (number of presences/number of samples). To also compare to the general Thai microbiome, 8 previously published 16s rRNA of microbial community Thai datasets were downloaded and analyzed (40,57–63). Within the derived studies, samples from non-disease (healthy control) individuals, regardless of age and sex, were included prior to any intervention, as part of the derived studies. The 16S analysis was performed as above, with minimal adaptations when required to fit the specifics of the individual dataset. Detailed methodological information for the additional datasets is provided in Supplementary Table 5. For more details see also the extended method section.

#### Differential abundance analysis

The differential abundance analysis was conducted by using ANCOM-BC2 with Pairwise Comparison mode (64). Analysis was performed at both the ASV level and genus level where taxonomic clustering and count summation were expected. Taxa with less than 10% prevalence and samples with a read depth below 1,000 were filtered out. To exclude potential confounding factors to the microbial composition, locality, sex, age, and BMI were included as fixed effects in the linear model. After getting the result, only microbes that are significantly different in at least one comparison (Phatthalung vs. Bangkok, Tak vs. Bangkok, and Tak vs. Phatthalung) were retained for further visualization and interpretation. For more details see the extended method section.

#### Lifetyle analysis

The lifestyle features with a high proportion of zeros (>50%) or missing values (>50%) were excluded. Pearson correlation analysis was then performed on the remaining features using the compositions v2.0.8 (56). During this step, features with zero variance across samples or those highly correlated with locality (Pearson correlation coefficient > 0.5) were removed. In addition, non-lifestyle features, including sex, age, height_cm, bmi, calprotectin, chromagranin, igm, iga, admixture, PRS_IBD, were manually removed. The remaining features were used in further analysis.

The filtered features were used to compute a lifestyle dissimilarity matrix. As the dataset contained mixed data types, Gower’s distance was calculated between samples using the cluster v2.1.6, followed by PCoA with the ape package v5.8 (55). To identify lifestyle features strongly associated with the PCoA ordination, lifestyle features were correlated with the first two axes of the Gower PCoA using Pearson correlation coefficients. Each feature was represented as a vector originating from the PCoA centroid (0,0), with vector coordinates corresponding to its correlations with PC1 and PC2. Vector length was calculated as the Euclidean distance from the centroid and used as a measure of the overall strength of association between each lifestyle feature and the ordination.

The importance of lifestyle features for distinguishing localities was assessed using a random forest model implemented in the randomForest package v4.7.1.2. Missing values in the lifestyle features were first imputed using the missForest package v1.6.1. The imputed dataset was then used to train the Random Forest classifier, and feature importance scores were extracted from the fitted model to identify lifestyle features contributing most to the classification of localities.

Further, to assess the association between lifestyle and microbial composition, a Mantel test was performed, correlating the β-diversity (Bray-Curtis, weighted UniFrac, and unweighted UniFrac) with the lifestyle Gower’s distance matrix using the vegan package v2.6.8 (47). For more details see the extended method section.

#### Host features and microbial association

Subsequently, lifestyle and non-lifestyle features (or host features) were examined for associations with microbial community α-diversity metrics (richness, evenness, Shannon, and Simpson indices) and phylogenetic diversity metrics (Faith’s PD and PSV). As the influence of lifestyle on diversity indices may depend on baseline microbiome differences across populations, analyses were conducted separately for each population.

For each dependent variable (diversity index), we performed feature selection to identify the subset of host features that best predicted the index. LASSO regression (glmnet v4.1.8 (65)) was applied with 10-fold cross-validation to determine the optimal λ, followed by fitting the LASSO model with this λ, repeated 400 times. Features selected in more than 80% of runs were retained. These selected features were then used to fit individual linear regression models for each diversity index using the lm() function in stats v4.4.1. This step was done to derive normal coefficient (not shrunk by LASSO) and statistical inference.

The same approach was applied to individual microbial ASVs to assess associations between the host features and ASV-level CLR-tranformed abundances. For more details see the extended method section.

#### Data handling and visualization

All plottings, and data manipulations were conducted in R v4.4.1. Taxonomic information, count data, and metadata were organized and managed using phyloseq v1.50.0 (54). General data handling and text manipulation were performed with tidyverse v2.0.0 (66), broom v1.0.7 (67), and stringr v1.5.1 (68). A map of study locations was created using sf v1.0.21 and rnaturalearth v1.1.0. Data visualizations were created using ggplot2 v3.5.2 (69), cowplot v1.1.3, and ggvenn v0.1.10. Final composite figures were assembled in Inkscape v1.4.2.

## Results

### Differences in the sample structure, demography and local factors

To characterize the microbiomes associated with different lifestyles, we isolated DNA from feces of 106 participants including foragers from the Phatthalung province (29 participants) and farmers from the Tak province (30 participants) and urban individuals from Bangkok (47 participants) (Figure 1A). Main geographic differences exist in the climate and population densities of the communities: Bangkok has the highest annual mean temperature (28.4 °C) and the lowest precipitation rate (1,358 mm), but by far the highest population density of all sampling sites (Figure 1B-D). The sampling was restricted to a few communities with heterogeneous demography, including age differences (slightly older in Phatthalung), BMI (slightly higher in Tak), and sex (more female participants in Bangkok residents, more male participants in Phatthalung) (Figure 1E-G). Furthermore, other differences existed as in the availability of electricity, the water sources, and the frequency of C-sections (Supplementary Figure 1 and Supplementary Table 1).

**Figure 1:**
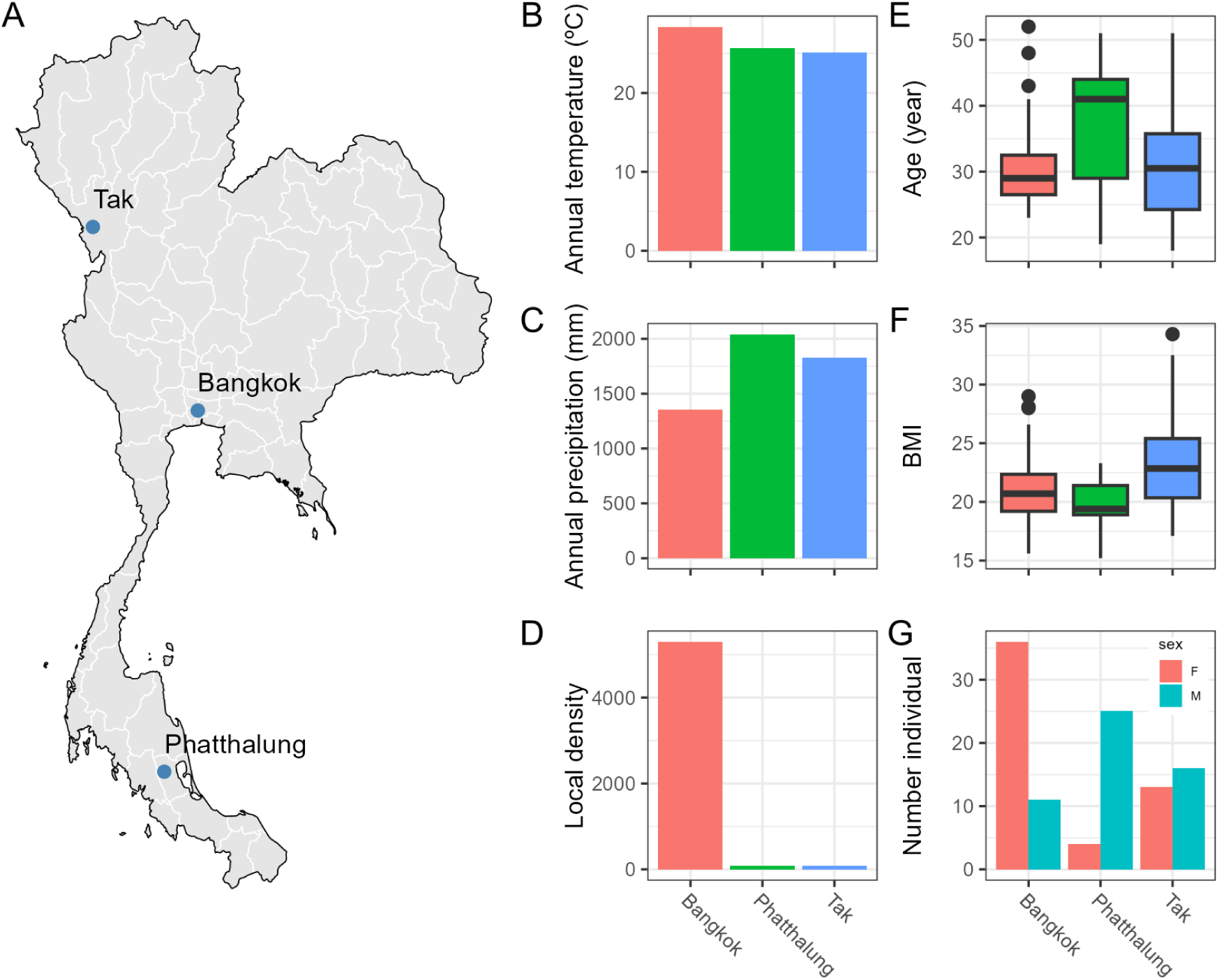
Sampling sites and main differences in demography and climate between sampled communities. Different sampling areas: Bangkok, Tak, and Phatthalung provinces in Thailand for urban, pre-industrialized farming, and foraging populations, respectively (A). Communities showed systematic differences in annual mean temperature (B), annual precipitation (C), and population density (D). Participants from different sampling sites differed in age (E), BMI (F), and sex distribution (G).

### Structure of microbial communities across populations

We used amplicon sequencing to describe microbial taxonomic diversity in the communities and found similar microbial communities on the phylum level mainly consisting of Bateroidota, Firmicutes, and Proteobacteria. Major differences existed in the expansion of Actinobacteriota and Fusobacteriota in the Bangkok cohort, an increase of the relative abundance of Firmicutes and Spirochaetota in Phatthalung, and a larger fraction of Bacteroidota in Tak cohort (Figure 2A). At the family level these phylum changes are mainly driven by the higher abundances in Bacteroidaceae and Bifidobacteriaceae (Bangkok), the increase of Oscillospiraceae, the Clostridia of the Methylpentosum group (both families Phatthalung), and the Prevotellaceae (Tak) (Figure 2B). The microbiomes of the different populations showed substantial overlap at genus level, with 137 of 236 genera (58%) shared across all groups. Further, the urban cohort from Bangkok displayed the highest specificity, with 36 (15%) genera unique to this population, followed by Phatthalung (13) and Tak samples (10) (Figure 2C and Supplementary Table 6).

**Figure 2:**
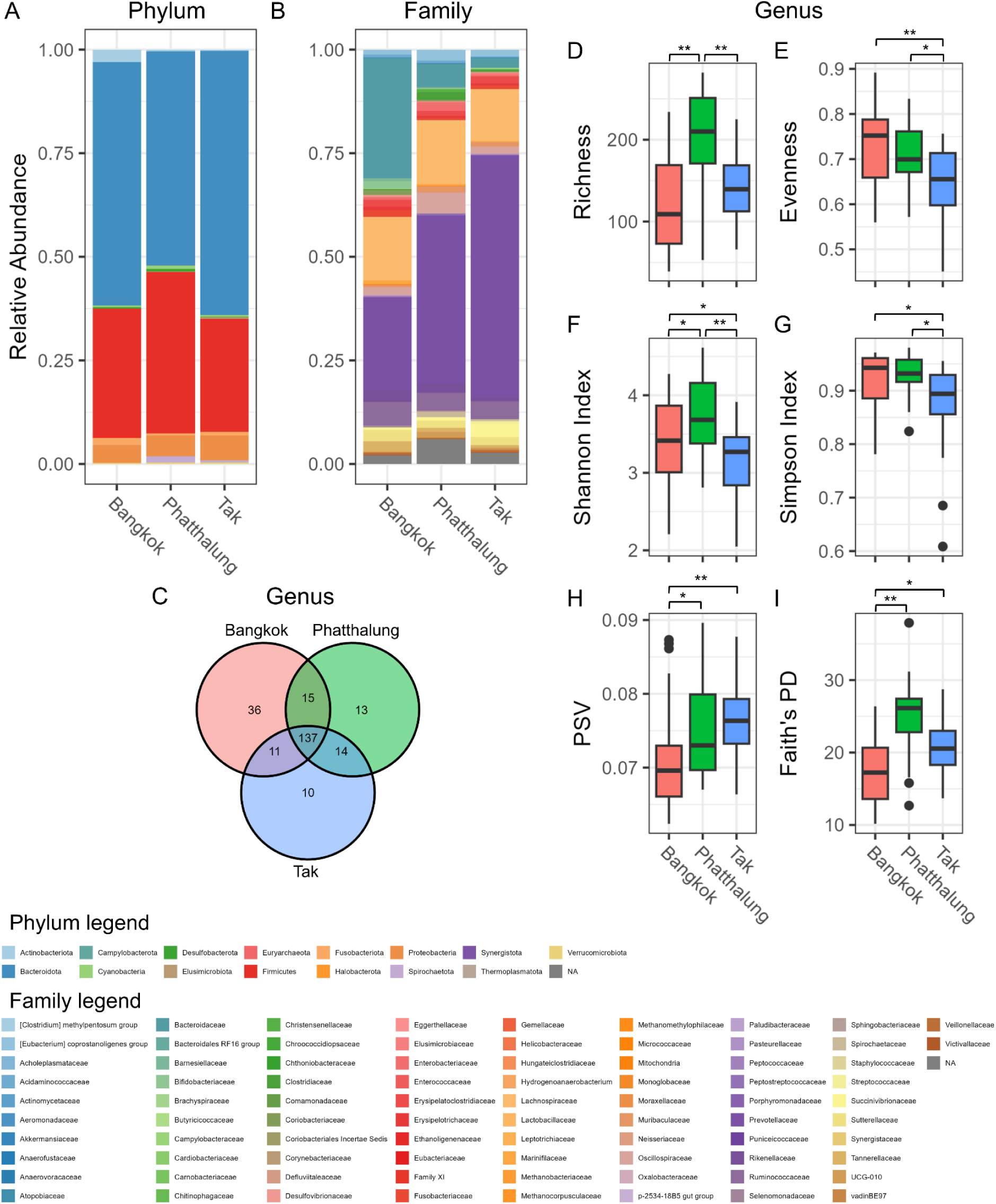
Microbial community overview and diversity index comparisons between populations. Overview of relative abundances on the phylum (A) and the family (B) level of detected species in the amplicon sequencing. Venn diagram shows the incidence of genera which are unique to or are shared between the three sampled populations (C). The diversity measures of Richness (D), Evenness (E), Shannon index (F), Simpson index (G), phylogenetic species variability (PSV) (H), and Faith’s phylogenetic diversity (Faith’s PD) (I) in Bangkok, Tak, and Phatthalung samples. Statistical significance was determined using a pairwise Wilcoxon test and significance levels are indicated as follows: * p < 0.05; ** p < 0.01.

We compared the three sampling sites in different α-diversity measures (Supplementary Table 7). Richness was highest in Phatthalung samples while evenness was highest in Bangkok samples (Figure 2D, E). This was reflected in Shannon (emphasizing richness) and Simpson index (emphasizing evenness), where diversity of Tak and Bangkok samples were highest, respectively (Figure 2F, G). When considering phylogenetic diversity, Bangkok samples consistently showed lowest diversity in phylogenetic species variability (PSV, Figure 2H) and Faith’s phylogenetic diversity (Faith’s PD, Figure 2I). Taken together, this indicates that Bangkok samples are characterized by a fewer and phylogenetically more closely related microbial species than Tak and Phatthalung samples.

We further examined the phylogenetic association and prevalence of identified microbes across populations (after rarefraction analysis, Supplementary Figure 2). The analysis revealed that although each population harbored area-specific genera (Figure 2C), the majority of highly prevalent microbes were shared across all three populations (Figure 3A, Supplementary Table 8). Among the population-specific taxa, the samples from Bangkok individuals exhibited the largest number of unique genera, mainly from the phyla Firmicutes and Actinobacteriota. In contrast, unique genera in the Tak samples were primarily from Proteobacteria, while those in the Phatthalung samples were distributed across smaller clades such as Thermoplasmatota, Halobacterota, and Verrucomicrobiota. Overall, most microbes were present in all populations, followed by population-specific taxa, and then those shared between two populations (Figure 3B, C). This pattern suggests highly overlapping microbiomes in the Thai population that persists despite differences in genetic background, geography and lifestyle. When we correlated genus prevalences of the different populations to each other, we observed the expected higher similarity of Phatthalung to Tak prevalences (r = 0.88), and Bangkok showing the smallest correlation coefficients to the other two data sets (r = 0.68 to Phatthalung and r = 0.80 to Tak, Figure 3B, Supplementary Table 9). Furthermore, microbial genera in the samples from Bangkok individuals exhibited the lowest prevalence across individuals, indicating greater heterogeneity within the population (Figure 3D), whereas the Tak and Phatthalung samples showed more homogeneous communities. Finally, we observed that genera with higher prevalence were more likely to be shared across populations (Figure 3E), highlighting that the shared taxa commonly detected in all populations also tend to be consistently found within individuals. To corroborate this analysis we gathered microbial datasets from six other locations (Buriram, Chachoengsoa, Chiang Mai, Chiang Rai, Maha Sarakham, Nakhon Phatom, Supplementary Figure 3, Supplementary Table 5) (40,57–63) and compared genus prevalence to this study (Figure 3A, Supplementary Table 8). The additional datasets were derived from both urban and rural populations and showed distinct patterns in their β-diversity ordination (Supplementary Figure 3), indicating a broad coverage of diverse Thai microbiomes. The prevalence analysis of these additional datasets confirmed the previous findings from the GMbC datasets. Microbial genera that were highly prevalent in one Thai population tended to also be highly prevalent in other populations (Figure 3A, B). These findings suggest the presence of a common set of highly prevalent microbial genera that are consistently observed across Thai populations, despite differences in lifestyle, and geographic location.

**Figure 3:**
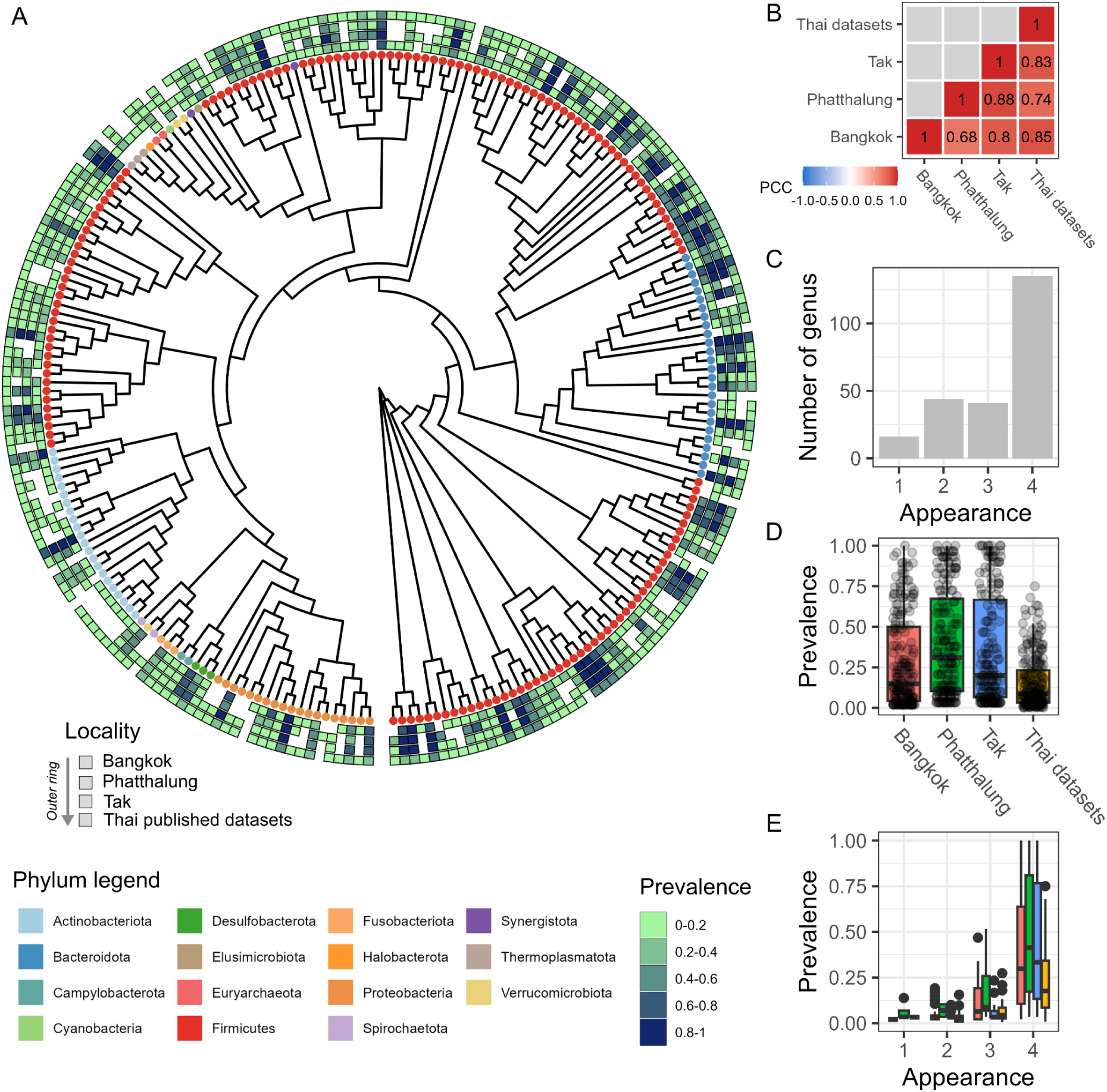
Phylogenetic tree and prevalence of genera across populations. Circular phylogenetic tree generated from genera representative amplicon sequences that tips of the tree represent microbial phyla (A). Heatmaps in the tree represent prevalence (frequency of a genus found in a population) of microbial genera found in each population and other Thai microbiome dataset. Pearson correlation coefficients (PCCs) of microbial genus prevalence between populations and the Thai microbiome dataset (B). Appearance (a genus found at least once within a population) of microbial genera (C). Prevalence of microbial genera in the different populations and the Thai dataset (D). Relationship of prevalence and appearance of microbial genera in different populations, Bangkok (red), Phatthalung (green), and Tak (blue), and the Thai dataset (orange) (E). In Figures A–E, the Thai datasets group included only genera that were recorded in this study (Bangkok, Phatthalung, and Tak).

### β-diversity and differential microbial abundance analysis in different populations

To further understand how microbial communities differ between populations, we calculated β-diversity measures (Aitchison and unweighted UniFrac) for all samples, tested for significant influence of the sampling location, and visualized the results with an ordination analysis (Figure 4A-B and Supplementary Figure 4). Overall, the microbiome derived from the Bangkok individuals showed strong separation from the other populations in all dissimilarity measurements, while foragers from the Phatthalung population and traditional farmers from the Tak population showed more overlap in Bray-Curtis and weighted UniFrac (Supplementary Figure 4). Here, Aitchison and unweighted UniFrac performed better in separating the microbial communities (Figure 4A-B). The Bray-Curtis, Aitchison, and weighted UniFrac measures showed significant differences in the β-dispersion between populations (ANOVA, p < 0.05), suggesting different spread of data between groups (Supplementary Table 11). However this was not observed in the unweighted UniFrac measure (p = 0.0658). 15.46% variance in dissimilarities for unweighted UniFrac were significantly explained by the three different sampled locations, with all three populations differing significantly in β-diversity measures (PERMANOVA, p = 9.999e-05) (Figure 4B and Supplementary Table 12). Pairwise comparisons of the unweighted UniFrac measure between populations were significantly different in all pairs (pairwise PERMANOVA, p adjusted = 0.0003) (Supplementary Table 13). Although β-dispersion in the other β-diversity measures were significantly different, we also performed pairwise comparisons on Bray-Curtis, Aitchison, and weighted UniFrac measures. The analysis results in all measures and population pairs were significantly different as shown in Supplementary Table 13.

**Figure 4:**
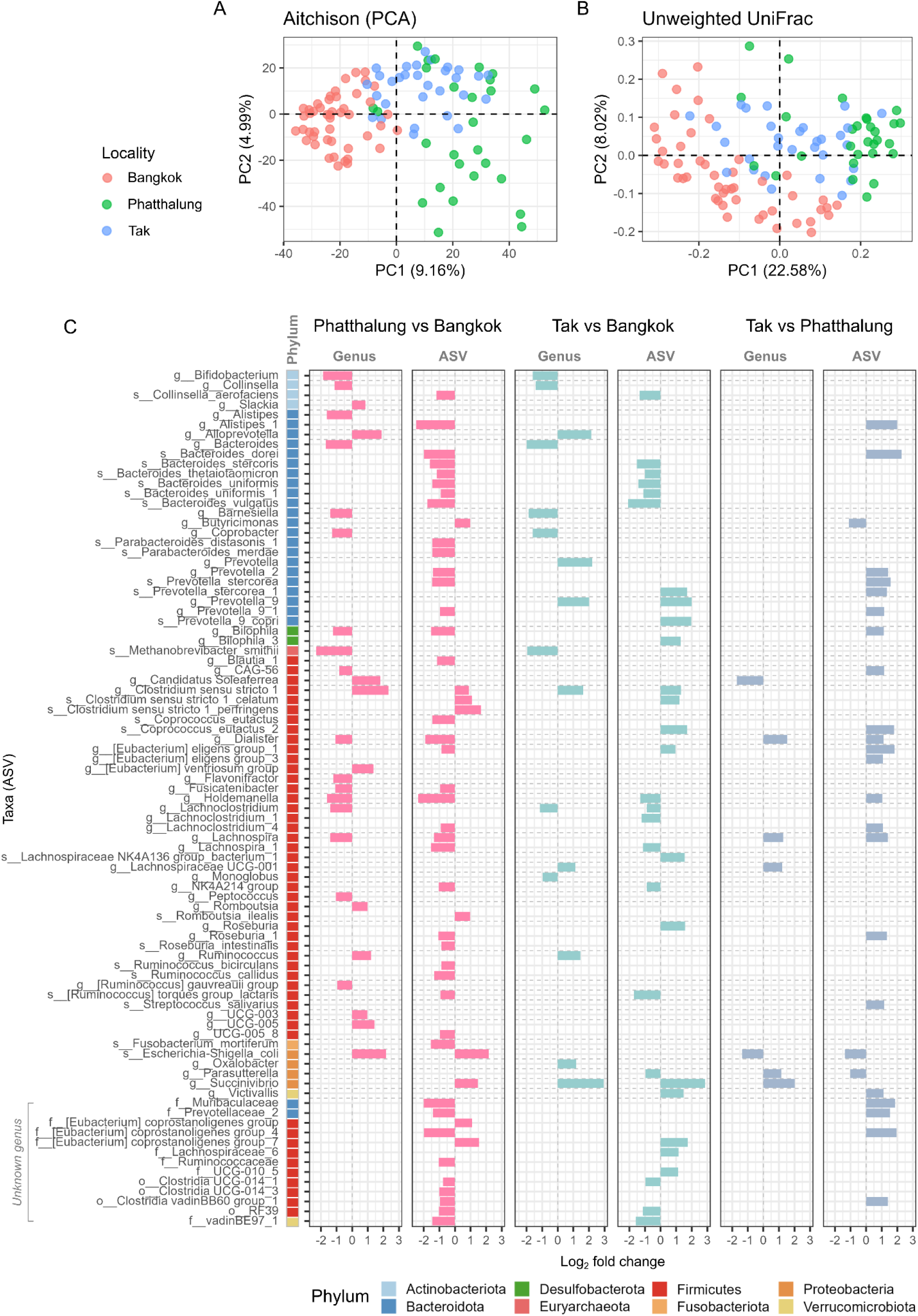
Ordination plots for β-diversity measures and pairwise differential microbial abundance analysis results. Principal component analysis (PCA) of Euclidean distance, Aitchison (A), Principal coordination analysis (PCoA) of unweighted UniFrac dissimilarity (B). Log_2_ fold change of microbial genera and ASVs comparing between pairs of populations, Phatthalung vs. Bangkok, Tak vs. Bangkok, and Tak vs. Phatthalung. The left color annotation indicates the phylum of each ASV, and horizontal dashed lines separate microbial genera.

We quantified differences of microbial communities in the three sampling sites on the genus level using ANCOM-BC (correcting for sex, age, and BMI - compare Figure 1). The highest number of differentially abundant taxa was identified in the comparison between Bangkok and Phatthalung samples (27 genera), followed by the comparison between Bangkok and Tak samples (16 genera). Least differential abundant genera were identified comparing samples from Tak and Phatthalung (7 genera) (Supplementary Table 14). The number of differentially abundant genera corresponded well to the analysis of the β-diversity, with microbial communities from Bangkok individuals being more dissimilar to those from Tak and Phatthalung individuals. The microbiomes from Bangkok individuals were significantly enriched in *Methanobrevibacter*, *Bacteroides*, *Bifidobacteria*, *Barnesiella*, *Bilophila*, *Coprobacter*, and *Lachnoclostridium*, while *Alloprevotella* and *Clostridium sensu stricto 1* were depleted comparing to Phatthalung and Tak individuals (Figure 4C). The *Escherichia*-*Shigella* and *Candidatus Soleaferrea* genera were specific for communities from Phatthalung and, in comparison to Bangkok and Tak, Phatthalung individuals showed lower abundances of both *Dialister* and *Lachnospira* (Figure 4C), which reportedly are associated to western and vegetable diets, respectively (70,71). Higher *Prevotella* abundance was identified comparing microbiomes from Tak and Bangkok individuals (Figure 4C). The samples from Tak individuals also showed higher *Succinivibrio* and *Lachnospiraceae* UCG-001 than the Bangkok and Phatthalung populations (Figure 4C). The genera specific for microbial communities from Bangkok are described in the literature as fermenters of diverse carbohydrates and polysaccharides (72–74), while the Phatthalung-specific genera are rarely found in healthy industrialized populations (75,76). Hence, we hypothesized that dietary and lifestyle habits will have a significant influence on the bacterial community composition.

We further analyzed differential ASV between the three populations and found similar results as in the genus-level analysis. Most ASVs differed in abundance between Bangkok and Phatthalung samples (49 ASVs), followed by the comparison of Bangkok and Tak (31 ASVs), and Tak and Phatthalung (25 ASVs) (Supplementary Table 15). Strikingly we found an enrichment of *E. coli/Shigella* and *Butyricomonas* ASVs in the Phatthalung samples. Bangkok samples showed consistent enrichment for different *Bacteroides* species (*B. vulgatus, B. stercoris, B. uniformis, B. thetaiotaomicron*) while Tak samples showed increased abundances in *Prevotella* species (two unidentified *Prevotella* species, and *Prevotella stercorea*), which we already observed on the genus level. Furthermore, Tak samples showed consistently increased abundances of *Coprococcus eutactus*, *Roseburia sp., Victivallis sp.,* and *Eubacterium eligens*. Other consistent indicators species for Bangkok microbiomes were *Holdemanella sp*., *Ruminococcus torques* (group lactaris), *Collinsella aerofasciens*, Bacilli RF39, *Lachnospira sp*., Clostridia UCG-014, and an unidentified species of the Oscillospiraceae NK4A214 group.

### Microbiome-lifestyle associations

Along with the microbiome sample collection, we conducted a self-reporting survey on medical treatments, food frequency, and other potential influencing factors for microbial community shaping. Firstly, we examined the associations of sampling locality with lifestyle-related features (excluding non-lifestyle-related variables; see Methods) to determine whether the populations differed in their lifestyles and which features contributed most to distinguishing the populations. We performed a PCoA analysis on the Gower’s distance based on the lifestyle features. The three populations do separate clearly in the ordination plot and the most important differences are determined by dietary habits (Supplementary Table 16), like the consumption of wild mammals (Phatthalung), avocado, ginger and cauliflower (Tak) or processed cereals, chocolate, butter and nuts (Bangkok, Figure 5A). A similar set of lifestyle features was identified, by a Random Forest classifier, as important for distinguishing the populations, providing consistent evidence across the two analytical approaches (Figure 5C, Supplementary Table 17). Overall, the results demonstrated lifestyle differences between studied populations.

**Figure 5.**
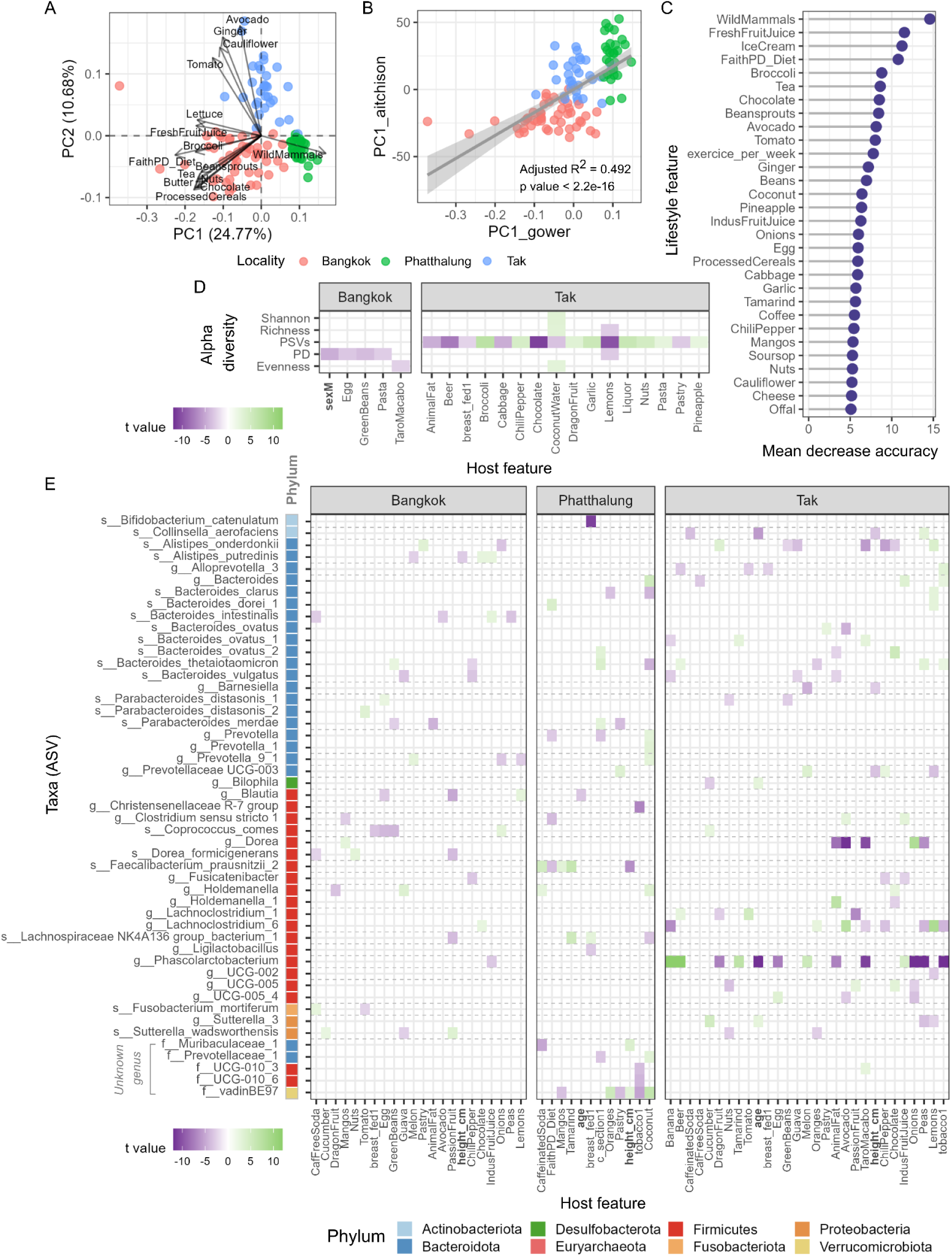
Lifestyle features differ across localities and are associated with variation in microbial diversity and community composition. (A) Principal coordinate analysis (PCoA) based on Gower’s distance of host lifestyle features across localities. (B) Association between the first principal component of Aitchison distance and Gower’s distance. (C) Top 30 mean decrease in accuracy of lifestyle features for predicting locality using random forest models. LASSO regression was used to identify lifestyle and genetic variables (host features) associated with variation in diversity measures and microbial abundance. Selected features for each diversity measure and ASV taxa abundance were subsequently fitted to multivariable linear models. Significant features identified by the multivariable linear models are shown for diversity (D) and microbial abundance (E) (adjusted p value < 0.05). Bolded features in the y-axis indicate non-lifestyle features. The 20 ASVs with the highest adjusted R² values for each locality and host features that were significantly associated with microbial abundance in more than three models are shown. Colored tiles indicate the effect size (t-value) of significant associations. The left color annotation indicates the phylum of each ASV, while horizontal dashed lines delineate different microbial genera.

Secondly, we investigated the relationship between overall lifestyle and microbial community structure. To assess whether differences in lifestyle were reflected in differences in microbial community composition, we examined the correlation between the first component of the microbial composition Aitchison-PCA and the first component of the PCoA based on Gower’s distance for the lifestyle features. We observed a strong correlation between the two ordination axes (R² = 0.492, p ≤ 2.2 × 10⁻¹⁶; Figure 5B). Further, we directly correlated microbial composition dissimilatory matrices (Bray-Curtis, weighted UniFrac, and unweighted UniFrac) to lifestyle dissimilatory matrix (Gower’s distance). The result showed slight but statistically significant positive correlations, for Bray Curtis and Unweighted UniFrac matrices (r = 0.13-0.15, all p < 0.02, Mantel test; Supplementary Figure 5 and Supplementary Table 18), supporting that the overall lifestyle exerts an effect on community structure.

Thirdly, we examined the influence of host features (include lifestyle and non-lifestyle features) on diversity measures and individual microbial taxa abundances. We stratified the dataset by the individual populations (to avoid collinearity) and applied LASSO regression to select features which explain differences in dependent variables best (see Methods). Afterwards, we tested statistical association of the identified features to different dependent variables in a linear model (Supplementary Figure 6, Supplementary Table 19). For the diversity, the result showed that, in the Bangkok samples, PD was negatively associated with male sex, as well as egg, pasta, and green bean consumption, while evenness was identified as negatively associated with taro consumption (Figure 5D). Coconut water consumption was associated with an increase in Shannon index, evenness and richness in the Tak population, while PSV was decreased (Figure 5D). Richness and PD was decreased with lemon consumption in Tak samples. Most features were significantly associated with PSV changes in the Tak population: most vegetables/fruits (broccoli, chilli pepper, dragon fruit, garlic, pineapple) increased the PSV, while most processed and animal derived foods (animal fat, beer, chocolate, pastry) decreased PSV. For the Phatthalung samples, we found no significant association of the host features to the diversity measures.

Furthermore, associations between individual microbial abundances and host features varied markedly across populations, with the broadest range of dietary and lifestyle associations observed in Tak, followed by Bangkok and Phatthalung (Figure 5E, Supplementary Figure 6, Supplementary Table 20). In Tak samples, *Phascolarctobacterium* was positively associated with banana, beer, tamarind and melon consumption, but negatively associated with egg, animal fat, taro, onion and pea consumption and tobacco use. *Dorea* also showed negative associations with animal fat, avocado and taro consumption. Avocado consumption was also negatively associated with the abundances of *Bacteroides sp.*, *B. intestinalis*, and *B. ovatus* in Bangkok and Tak, respectively. Green bean consumption in Bangkok and pea consumption in Tak were associated with increased abundance of *B. thetaiotaomicron*, whereas guava consumption was associated with decreased abundance of *B. vulgatus*. Interestingly, the abundances of *Prevotella sp.*, *Prevotella_9 sp.*, and an ASV of the family Prevotellaceae were positively associated with coconut consumption in the Phatthalung population. Among the highlighted host traits, age was negatively associated with *Phascolarctobacterium* and *Collinsella aerofaciens* in Tak and with *Blautia* in Phatthalung. Height showed several negative associations in Tak samples: *Prevotella UCG-003*, *Barnsiella*, *Collinsieall aerofaciens*, while *Alistipes putridines* was negatively associated with height in the Bangkok population. In Phatthalung, height was negatively associated with a *Faecalibacterium prausnitzii* ASV. In general, the host-factor-ASV association showed no obvious pattern, indicating a population specific response of microbial compositions to differences in the lifestyle making it difficult to predict microbial communities from host factors alone.

## Discussion

In this study, we characterized the gut microbiota of Thai populations representing distinct lifestyles, urban population, pre-industrialized farmers, and rural foragers, originating from the Bangkok, Tak, and Phatthalung provinces, respectively (Figure 1A). To our knowledge, this is the first report describing the gut microbiomes together with lifestyle of individuals from these rural populations in Thailand. Our results showed distinct demographic and lifestyle patterns across the three populations (Figure 1B–G, Figure 4A, Supplementary Figure 1, and Supplementary Table 1). Demographically, populations from more industrialized settings had greater access to electricity, treated water, and both antibiotics and traditional medicines, compared with the more rural populations. Interestingly, across the Thai populations, vaginal delivery and breastfeeding remained relatively common, indicating similarities in these aspects of maternal and infant care despite differences in modernization and accessibility. Regarding lifestyle, the Bangkok population was characterized by a more modernized dietary pattern, including consumption of processed, high-sugar, and high-fat foods, whereas the Tak population showed greater reliance on plant-derived foods and the Phatthalung population reported a greater contribution of forest-derived foods (Figure 4A, 4C, and Supplementary Table 1). Together, these differences indicate variation in the degree of modernization and industrialization across the studied populations, consistent with their recognized dietary and subsistence practices and previous reports (33,36,40).

Analysis of microbial profiles revealed clear compositional differences among the three populations, with significant variation in both α-, phylogenetic- and β-diversity. We observed a higher richness and phylogenetic diversity in non-industrialized samples and distinct clustering in the β-diversity ordinations, compared to industrialized samples. Particularly, Faith’s PD and PSV indicated that the microbiomes from the rural samples exhibited higher phylogenetic heterogeneity compared to the urban samples. The loss or reduced persistence of certain microbial taxa in samples from Bangkok individuals is potentially driven by higher sanitation, increased antibiotic use, or industrialized lifestyle factors and may have narrowed the phylogenetic breadth of the microbiome (14,77). In this regard it is interesting to note that while the individual α- and phylogenetic diversity was lower in Bangkok samples compared to those from Phatthalung, the population wide diversity was higher, indicated by a greater number of unique genera and a wider spread of samples in the unweighted Unifrac ordination (Figure 2C, 4B). This most likely reflects the higher variability of lifestyle choices participants are able to make in urban and industrialized environments. When comparing microbial compositions of the three sampling sites, we generally find industrialized samples show lower abundances in Prevotellaceae and higher abundances in Bacteroidaceae which is in line with previous reports (8,9,25,78). Industrialized microbiomes have been described to be depleted in VANISH microbes (volatile and/or associated negatively with industrialized societies of humans) such as in microbial families of Prevotellaceae and Succinivibrionaceae (79–81), which we also confirm in our study.

Similar patterns have been observed across Southeast Asian populations. Specifically for traditional farmers of the Tak province it has been described that emigration to the US induces a loss of diversity and Bacteroides to Prevotellaceae replacement (57). Our study confirms these findings between Tak and Bangkok populations within Thailand pointing that industrialization is causing these changes. Conversely, it indicates that genetics have a smaller effect on microbial composition (as also described in Dwiyanto et al., 2021 and Seetaraso et al., 2025 (82,83)) and that diet and environment are the main drivers, at least for Thai individuals. Studies in Indonesia and Malaysia likewise identified urbanization-related changes alongside substantial variation between local communities (84,85). These findings indicate that urbanization-associated microbiome changes depend strongly on local diet, environment, and population history. Interestingly, Tee et al. (85) found increased microbial diversity with increased levels of helminth infections, showing that hygiene is a major contributor to microbiome community structure. Other studies showed that low carbohydrate diets are associated with the Prevotella-to-Bacteroides switch in western (86) and south east asian populations (87). Notably, a study in Vietnam compared rural and urban microbiomes and found increased α-diversity in the urban population and associated that to differences in dietary habits between the populations (88). While this looks contrary to our findings, it confirms the notion that diet has the strongest impact on the microbiome composition.

The pronounced differences in diversity measures in samples from Tak and Phatthalung individuals also suggest that they should not be simply grouped within the “non-industrial lifestyle” category. Samples from the Phatthalung individuals exhibited higher Bacteroidaceae but lower Prevotellaceae abundance compared to those from Tak, while the reverse was observed for Succinivibrionaceae. These observations contrast with previous reports (79,80) and highlight that microbiotas adjust to cultural shifts and support human (rapid) adaptations to new lifestyles (89,90). Further, we observed a gradient decline of transmission and persistence of certain phylogenetically distant microbes from foraging over pre-industrialized farming to an industrialized lifestyle. For instance, most genera of archaeal phyla such as Thermoplasmatota, and Halobacterota as well as single genera in the Verrucomicrobiota, Firmicutes, and Bacteriodota could be detected in Phatthalung individuals but were absent in the other populations (Figure 3A, Supplementary Table 8). On the other hand, we detected many genera unique to the urban populations (Bangkok, other Thai) belonging predominantly to the Actinobacteriota and Firmicutes phylum (Figure 2C, and 3A). Additionally, while higher PSV in microbiomes from Tak over Phatthalung individuals was observed (Figure 2H), we found a lower Faith’s PD in the Tak samples (Figure 2I). These results suggest that a limited set of microbial taxa dominate, indicating that the switch from a foraging to a farming to an industrialized lifestyle potentially prevents the proliferation of a broader range of microbial lineages.

Associations between lifestyle factors, and disease have become increasingly evident, with conditions such as obesity, type 2 diabetes, depression, and cancer occurring more frequently in industrialized populations (15,91–95). This study revealed population-specific microbial signatures (Figure 4C-E) that may correspond to lifestyle-associated diseases. For example, in Bangkok individuals, increased abundances of *Methanobrevibacter* and *Lachnoclostridium*, and *Collinsella* taxa are linked to increased body weight and obesity risk (96–98). However, the Bangkok individuals also exhibited a marked increase in *Bifidobacterium*, a genus that has been associated with reduced fat storage (98,99) and is considered health beneficial. The Tak individuals, on the other hand, showed enrichment in *Prevotella, Alloprevotella,* and *Lachnospiraceae* UCG-001, which has been associated with reduced obesity and diabetes risks (97,100,101). In contrast, the Phatthalung individuals showed reduced abundance of *Barnesellia* which has been associated with an increased diabetes risk (101,102). This finding aligns with a previous study that foraging communities may have limited tolerance to refined carbohydrates and face an elevated diabetes risk when adopting a Westernized diet (103), which might be partly driven by their microbiome composition. Furthermore, Phatthalung individuals showed increased abundances of *Slakia*, *Clostridium sensu stricto 1,* and *Lachnoclostridium* which have been associated with reduced cancer risk (104–106).

Overall lifestyle variation was highly correlated with microbial composition changes (Figure 5B). We found most associations of lifestyle to specific microbial genera in the samples from Tak individuals, followed by the samples from Bangkok and Phatthalung individuals (Figure 5B). This reflects on the one hand the level of dispersion in the β-diversity of the different populations and on the other hand the more diversity of lifestyle choices the participants are able to make. Hence, it directly links the environmental diversity to the microbial diversity. However, the increased dispersion in Bangkok and Tak samples does not translate to increased diversity on the individual level. In addition, among the selected host factors, dietary features were most frequently identified, highlighting the potentially predominant role of diet in reshaping the gut microbiome compared to other lifestyle factors. In Tak, *Phascolarctobacterium* sp. abundance was associated with several dietary habits. As this bacterium utilizes fermentation products such as succinate generated by other gut bacteria, these associations may reflect indirect dietary effects mediated by changes in the activity or abundance of succinate-producing bacteria (107–109). Similarly, we identified *Dorea sp.* in Tak individuals as dependent on dietary habits, which has been reported before (110). Moreover, the results suggest that microbial abundance responded in a population-specific manner to lifestyle influences given the high number of uniquely identified taxa-lifestyle associations. This may reflect differences in baseline microbial composition across populations, leading to distinct responses to lifestyle factors and matches dietary intervention studies in western populations, where response is often highly dependent on the resident microbiome (111–113). There were least microbiome-lifestyle associations in the Phatthalung population, probably reflecting the decreased variability of lifestyles and dietary habits in these communities and higher homogeneity of microbial community composition.

However, conclusions on these results are limited by the study design and sampling. For instance, we only consider taxonomic information on the 16S level, neither resolving strain levels nor functions of the identified bacteria. Furthermore, our study is limited to a rather small number of participants and only one representative location for foraging, farming and urban lifestyle. The study does not consider socioeconomic variables as host factors which might be an important influence on the microbiome composition (114,115). This is further complicated by minor, yet present differences in genetic background of our populations (35,38,39), which is not regarded in our analysis but potentially contributes to the observed differences in the microbial composition. Nonetheless, our results suggest that gut microbiomes from different populations respond differently to lifestyle factors, highlighting the importance of considering lifestyle-specific and population-specific contexts in microbiome research. These findings not only contribute to expanding the lifestyle-dependent global microbiome catalog but also provide a foundation for future studies aiming to connect lifestyle-associated microbial variation with health outcomes in underrepresented populations. Furthermore, we highlight the gradual shift in microbiomes from foraging over farming to urban populations, which emphasize the high interdependency between lifestyle (and diet) and the microbiome as well as the potential role of the microbiome to adapt to new environmental conditions. Finally, the study shows that the majority of microbial taxa is shared across the different populations which also contribute most to relative abundances in our analysis. This suggests that lifestyle alterations influence microbial composition to a limited extent, only a fraction of taxa respond by being gained or lost, while the majority persist.

## Data Availability

The 16S sequencing data will be made available online on the dbGaP server (Study ID: 38715; Accession: phs002235.v1.p1) upon publication of the article. Meta, lifestyle data was derived from (29). The GMbC participant metadata can be requested through the Global Microbiome Conservancy biobank (https://microbiomeconservancy.org/). All data and metadata are distributed via controlled-access systems in accordance with the original informed-consent provisions. Access to metadata requires a formal application and a data-access agreement. Count data and scripts for data analysis and visualization are provided via github.com/nsrinak/GMbC-Thai-population.

## Ethics approval

Research & ethics approvals were obtained from the MIT IRB (protocol #1612797956) and from the Ethics commission of the Medical Faculty of Kiel University (Studie D 511/24). Permits were also obtained in Thailand prior to the start of sample collection from the Human Research Ethics Committee of the Faculty of Medicine of Thammasat University No.1, Certificate of Approval 143/2019. All participants provided both oral and written informed consent in their native language to study personnel on site.

## Author contribution

NS - Formal analysis, Data Curation, Writing - Original Draft, Visualization

RV - Investigation, Resources, Data Curation

LMS - Resources, Data Curation

CC - Resources, Data Curation

CK - Funding acquisition

EJA - Funding acquisition

MG - Conceptualization, Resources, Data Curation, Project administration, Funding acquisition

VM - Investigation, Resources, Data Curation, Project administration,

MP - Conceptualization, Resources, Investigation, Project administration, Funding acquisition, Supervision

JT - Conceptualization, Writing - Original Draft, Supervision, Project administration, Funding acquisition

All authors - Writing - Review & Editing

## Acknowledgments

This work was supported by grants from the Center for Microbiome Informatics and Therapeutics at MIT and the Rasmussen Family Foundation to EJA, M.G. & M.P. We acknowledge support by the German Research Foundation within the framework of the Excellence Cluster “Precision Medicine in Chronic Inflammation” (project code EXC2167) and the research group miTarget (project code FOR5042), the project ExoMod (project code KA 3541/20) to C.K. and individual funding (project code TA1699.2) to J.T.. Moreover, we acknowledge support by the German Ministry for Education and Research within the scope of e:Med iTREAT (project code 01ZX1902A) to C.K.. M.P and M.G. received support from the Deutsche Forschungsgemeinschaft (DFG - German Science Foundation) within the Collaborative Research Center (CRC) 1182 on “The Origin and Function of Metaorganisms” (Project-ID 261376515 – SFB 1182, project C5.1 to M.G., project C5.2 to M.P.). We thank the Infectious Disease and Microbiome Program’s Microbial Omics Core at the Broad Institute of MIT and Harvard for their support on sequencing efforts. We thank Dr. Claudia Taubenheim for fruitful discussions and for carefully reading and editing the manuscript.

## Supplementary Data

Supplementary File 1: Extended methodology.

Supplementary Table 1: Summary of host features in each locality.

Supplementary Table 2: Scoring scheme of host features.

Supplementary Table 3: General statistic of 16s rRNA sequencing data.

Supplementary Table 4: Read count for each microbial taxa.

Supplementary Table 5: Previously published Thai microbiome datasets.

Supplementary Table 6: Rare genus found in different populations.

Supplementary Table 7: Pairwise comparison of diversity indexes between populations using Wilcoxon Ranked-Sum test.

Supplementary Table 8: Prevalence of microbial genera for each locality.

Supplementary Table 9: Pearson correlation of microbial prevalence between localities.

Supplementary Table 10: Prevalence of microbial species for each province in Thailand.

Supplementary Table 11: β-dispersion and comparison using ANOVA in different populations.

Supplementary Table 12: PERMANOVA analysis of different β-diversity indexes.

Supplementary Table 13: Pairwise PERMANOVA across different populations.

Supplementary Table 14: Pairwise differential abundance analysis at genus level using ANCOM-BC2 across different populations.

Supplementary Table 15: Pairwise differential abundance analysis at ASV level using ANCOM-BC2 across different populations.

Supplementary Table 16: Correlation vectors of lifestyle features for PC1 and PC2 in PCoA based on Gower’s distance.

Supplementary Table 17: Mean decrease of accuracy of lifestyle features to locality from random forest model analysis.

Supplementary Table 18: Mantel test between microbial composition and lifestyle across different populations.

Supplementary Table 19: Associations between host features and alpha diversities based on linear regression.

Supplementary Table 20: Associations between host features and microbial ASV abundance based on linear regression.

## Supplementary Figure

**Supplementary Figure 1:**
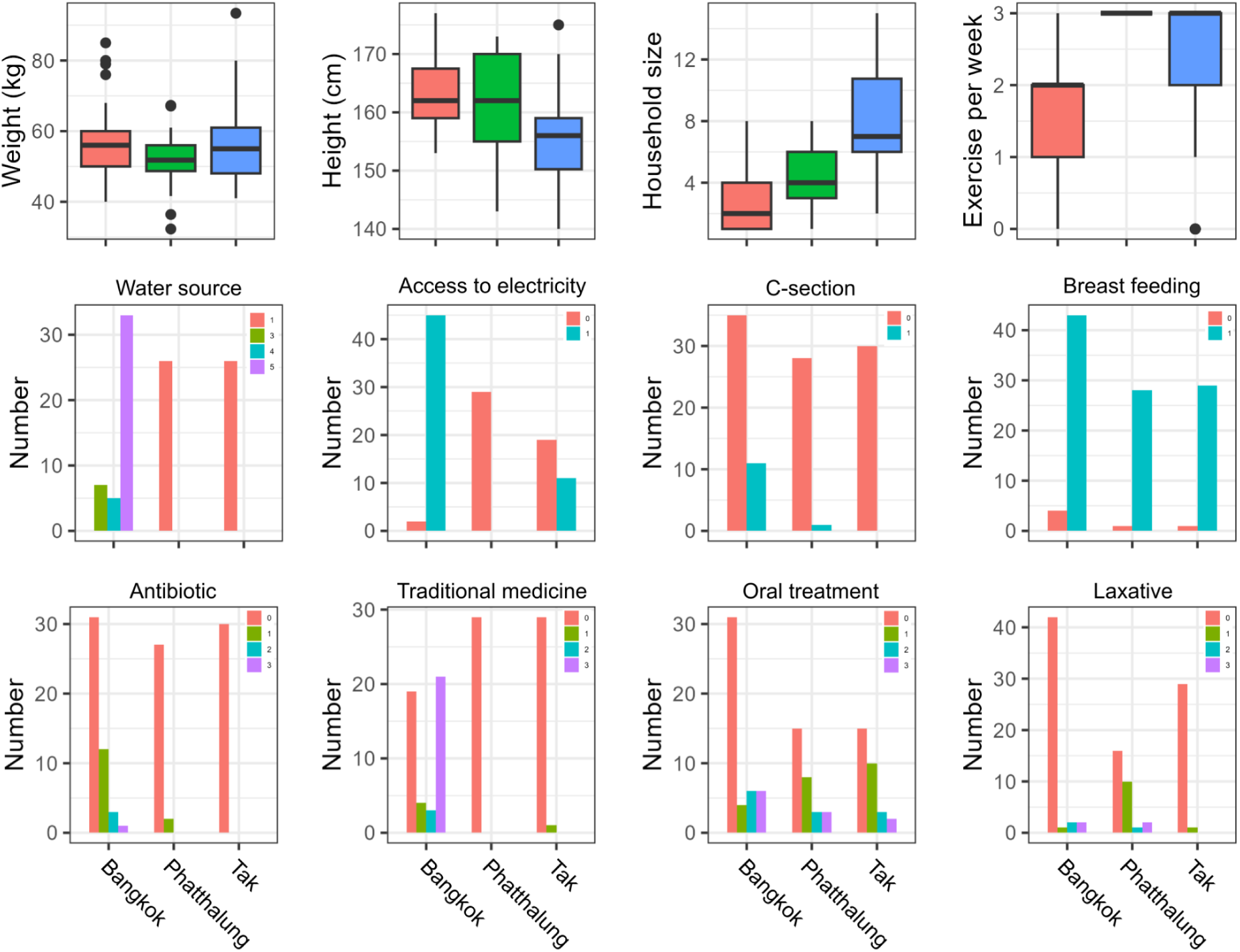
Differences in demography between sampled communities. For water source, scores of 1–5 indicate increasing levels of water treatment, ranging from untreated and unfiltered to untreated and filtered, and treated and filtered water. For access to electricity, C-section, and breastfeeding, scores of 0 and 1 indicate “No” and “Yes,” respectively. For antibiotic use, traditional medicine use, oral treatment, and laxative use, scores of 0–3 indicate the following time intervals: more than 6 months ago or never, within the past 6 months, within the past month, and within the past week, respectively. The complete scoring scheme and interpretation of all variables are provided in Supplementary Table 1.

**Supplementary Figure 2:**
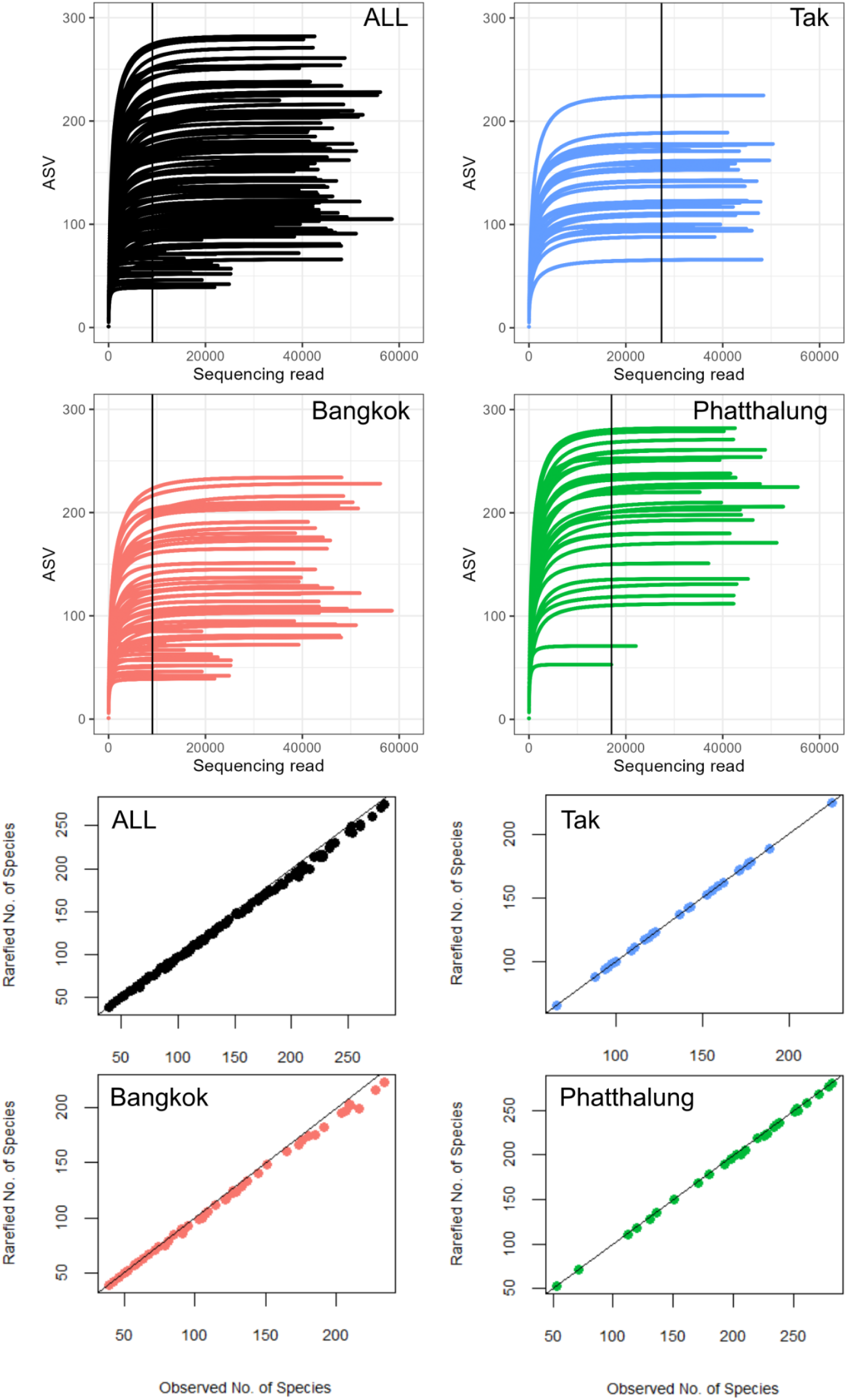
Rarefaction curves with horizontal lines of minimum sequencing depths and dot plots of observed species against number species from rarefied samples.

**Supplementary Figure 3:**
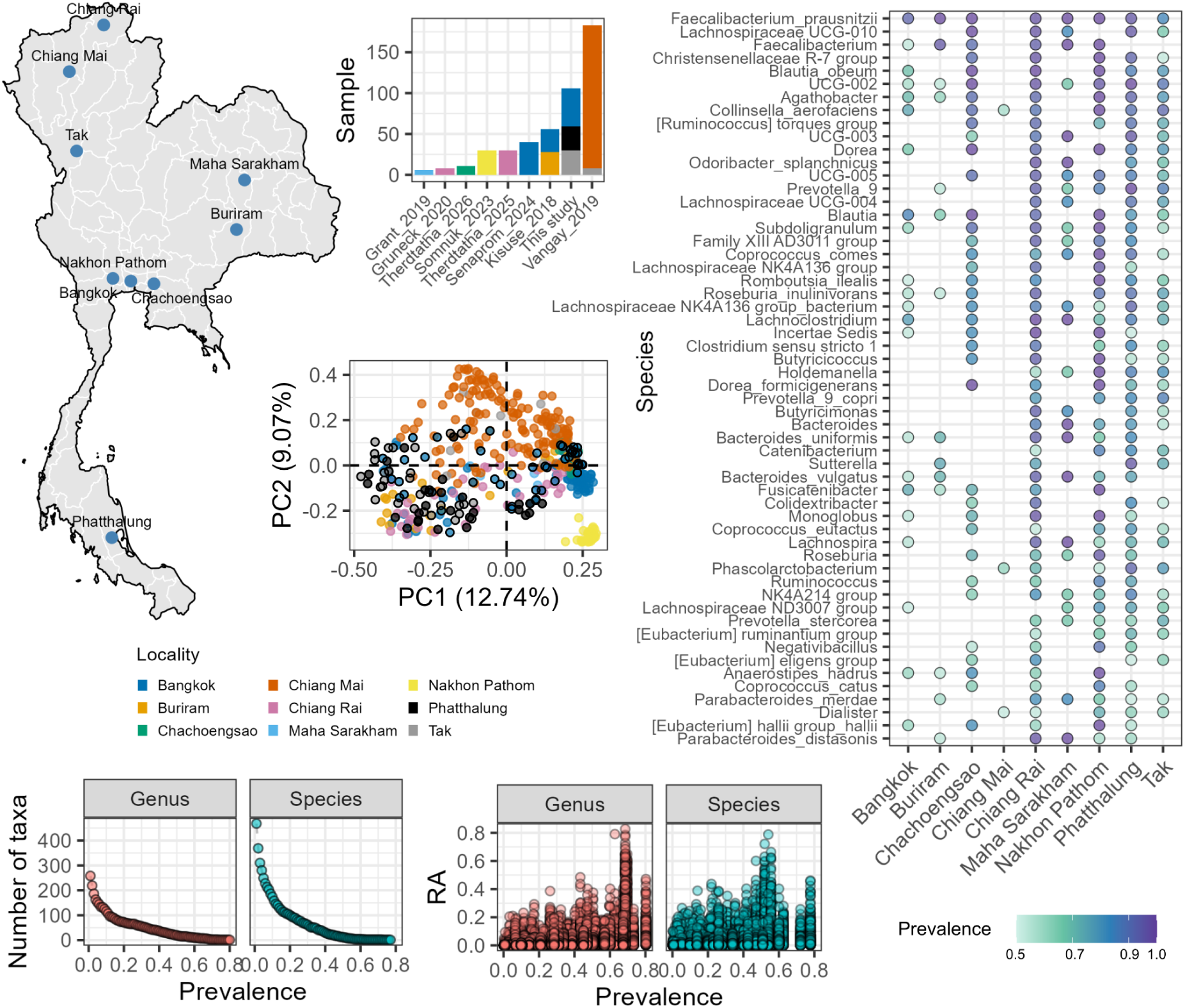
Microbial species prevalence across previously published Thai 16S rRNA gene sequencing datasets. Map of Thailand showing the locations of datasets, including those from this study, covering nine provinces (A). Number of samples from each study, grouped by locality (B). Principal coordinate analysis (PCoA) based on Bray–Curtis dissimilarity of microbial community composition. Samples from this study are indicated by black outlines (C). Number of genera and species retained at different prevalence thresholds (D). Association between microbial relative abundance and prevalence at the genus and species levels (E). Bubble plot showing microbial species with a prevalence >0.5 in more than three locations (F). The microbial species prevalence for each location can be found in Supplementary Table 10.

**Supplementary Figure 4:**
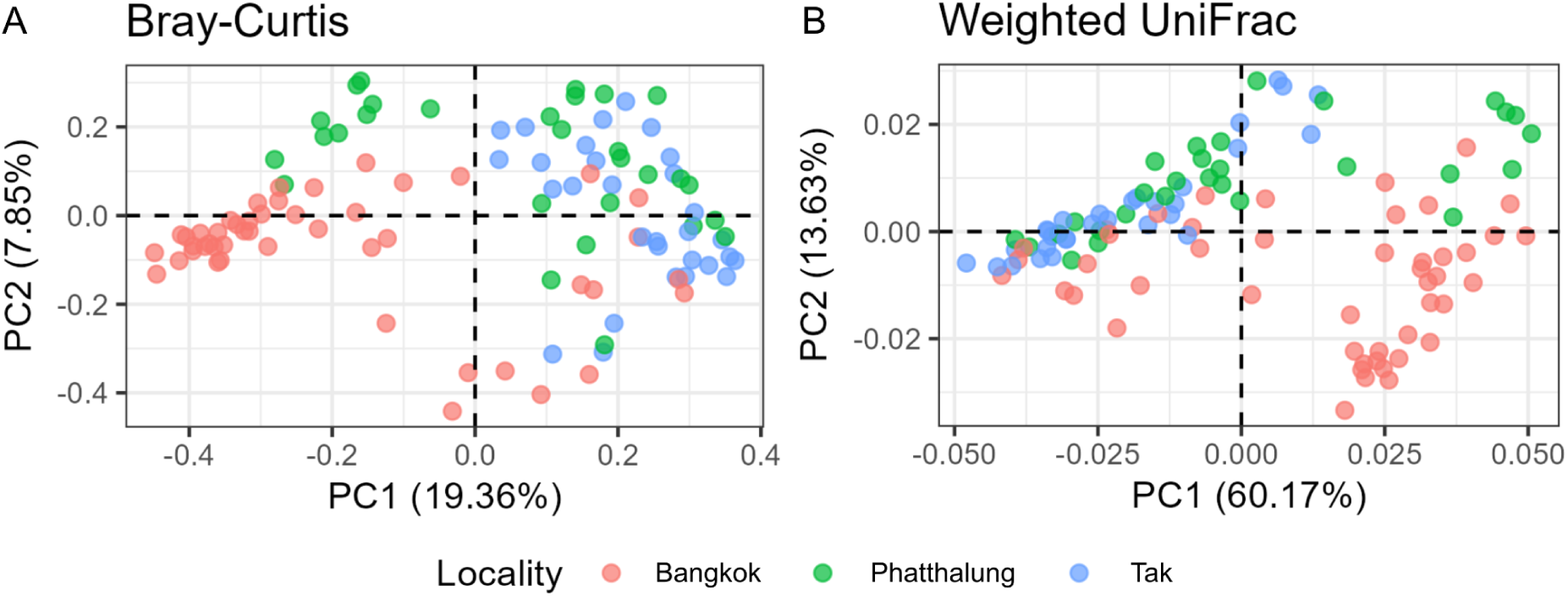
Ordination plots for β-diversity measures. Principal coordination analysis (PCoA) of Bray-Curtis (A), and weighted UniFrac dissimilarity measures (B).

**Supplementary Figure 5:**
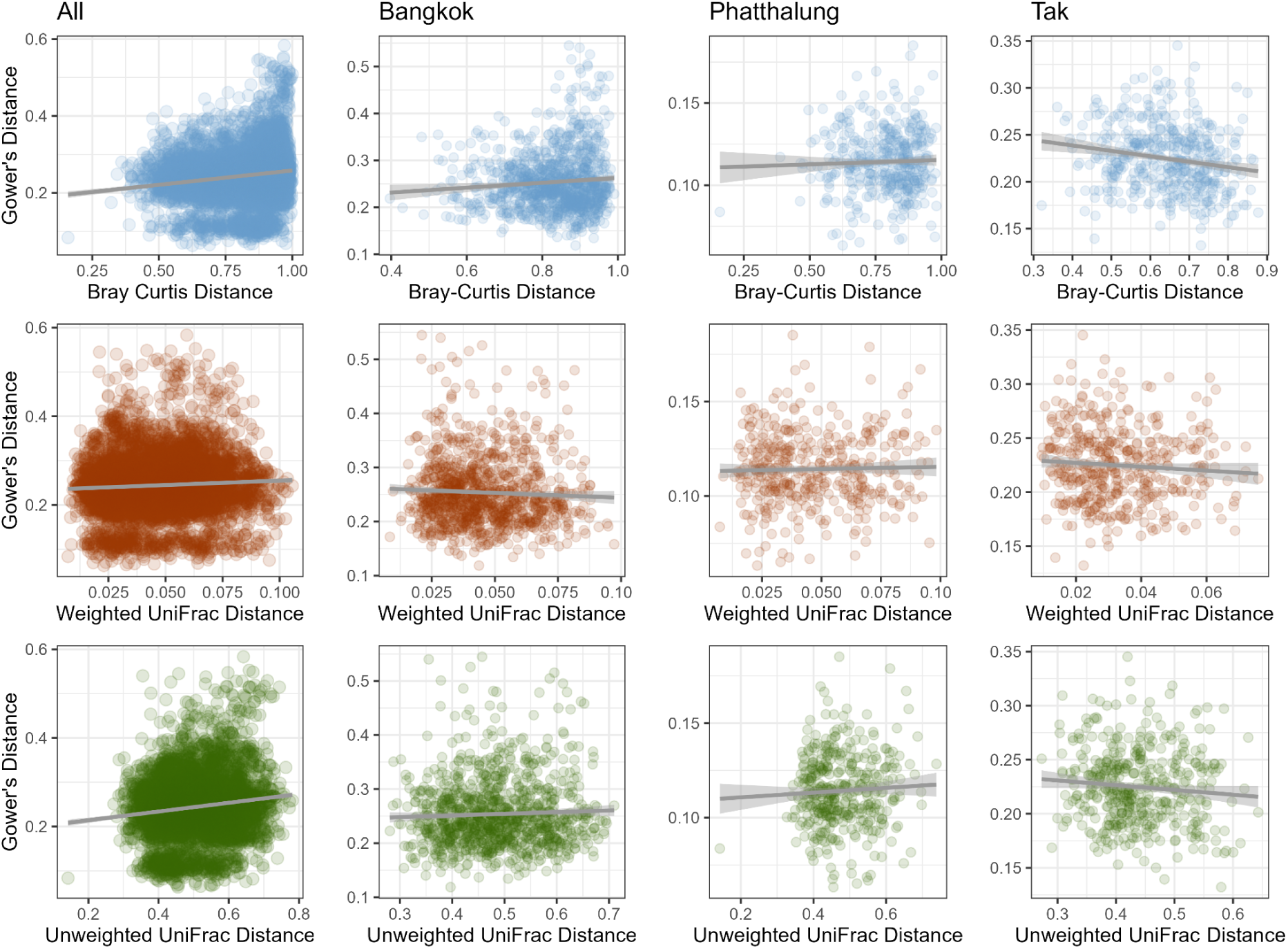
Associations of lifestyle distance and microbial β-diversity indexes. Associations of the Gower’s distance of host lifestyle to Bray-Curtis, unweighted UniFrac, and weighted UniFrac distances of all populations combined and separately for each population: Bangkok, Phatthalung, and Tak.

**Supplementary Figure 6:**
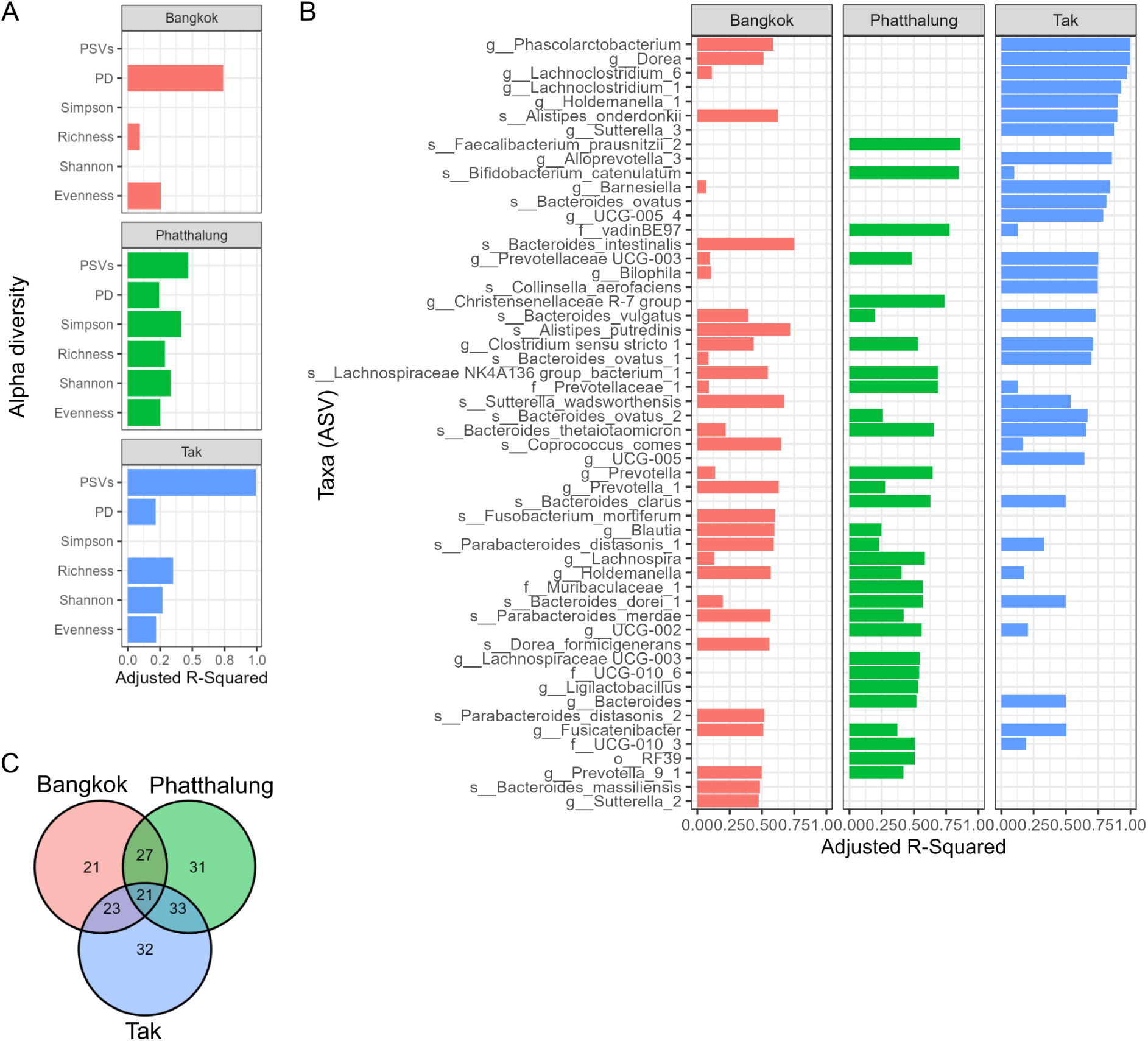
Lifestyle differences significantly associate with changes in different microbial composition. LASSO regression was used to identify important lifestyle and genetic variables (host features) which explain differences in the microbial diversity and individual abundance. The features for each microbial feature were later fitted to multivariate linear models. The barplot indicates significant adjusted R² from the linear model after the final chosen LASSO model for each different diversity index (A) and microbial ASV (B) (adjusted p value < 0.05). For B, ASV taxa with top 20 adjusted R² for each location were selected to visualize. Venn diagram of significantly predicted ASV across populations (C). All adjusted R² from microbial abundance and host feature association can be found in Supplementary Table 20.

