## Extended Methods for "Variation in gut microbiome diversity and structure across host lifestyles in Thailand"

#### 16S rRNA identification

Paired-end 16S rRNA gene sequencing reads were processed using a quality-control, denoising, and taxonomic-classification workflow. Paired-end sequencing generates two reads for each DNA fragment—one from each end—which can improve sequence accuracy and coverage of the targeted 16S rRNA gene region.

Initial read quality was assessed using FastQC v0.12.1 (1). FastQC examines characteristics such as the quality score at each nucleotide position, read length, GC content, ambiguous bases, and the presence of overrepresented sequences or sequencing adapters. MultiQC v1.25.1 (2) was subsequently used to combine the individual FastQC reports into a single summary, allowing quality patterns and potential problems to be compared across all samples. The reads had a mean GC content of 51.92%, and samples contained an average of 46,934 reads. Most samples (197 of 212) displayed consistently high Phred quality scores. Phred scores express the probability that a nucleotide was called incorrectly; higher scores therefore indicate greater confidence in the reported nucleotide sequence.

The reads were then filtered and trimmed using the `filterAndTrim` function of the DADA2 package (3). During this step, `truncQ = 11` was used to truncate a read when a nucleotide with a quality score of 11 or lower was encountered. In addition, `maxEE = 2` excluded reads with more than two expected sequencing errors. Expected-error filtering considers the quality scores across the entire read and estimates the total number of incorrect nucleotide calls likely to be present. It is therefore generally more informative than filtering a read solely on the basis of its lowest individual quality score. These filtering steps reduce the contribution of unreliable sequences to downstream analyses.

The quality-filtered reads were subsequently processed with DADA2 to infer amplicon sequence variants (ASVs). DADA2 models the characteristic sequencing-error patterns in the dataset and uses this model to distinguish genuine biological sequences from errors introduced during sequencing. Unlike traditional operational taxonomic unit clustering, which groups sequences according to a predefined similarity threshold—commonly 97%—ASV inference can distinguish sequences that differ by a single nucleotide. This provides a higher-resolution and reproducible representation of the bacterial sequences detected in each sample. The resulting ASV table recorded the number of times each ASV occurred in every sample.

Taxonomy was assigned to the ASVs using a naive Bayesian classifier trained on the SILVA non-redundant 99% reference database, version 138.1 with species-level annotations (`SILVA nr99 v138.1_wSpecies`) (4). The classifier compares the nucleotide composition of each ASV with that of taxonomically annotated reference sequences and estimates its most likely classification at successive taxonomic ranks, such as phylum, class, order, family, genus, and, where sufficient sequence information is available, species. Because short 16S rRNA gene regions do not always contain enough variation to discriminate closely related species, species-level assignments were made only when supported by the reference sequence and the information contained in the amplified region.

FastQC and MultiQC were run from the Bash command line. Read filtering, ASV inference, construction of the ASV table, and taxonomic assignment were performed in R v4.4.1 using DADA2.

### Statistical analysis

#### α-diversity

Within-sample microbial diversity (α-diversity) was characterized using several complementary indices. Whereas taxonomic α-diversity describes the number and relative abundances of ASVs within a sample, phylogenetic α-diversity also considers the evolutionary relationships among those ASVs.

The Shannon and Simpson diversity indices were calculated using the vegan package v2.6.8 (5). The Shannon index accounts for both richness—the number of ASVs detected—and evenness—the similarity of their relative abundances. It increases when a sample contains more ASVs or when their abundances are distributed more evenly. The Shannon index was calculated as

$$H' = -\sum_{i=1}^{S} p_i \ln(p_i),$$

where (*S*) is the number of ASVs in a sample and *(p_i)* is the relative abundance of ASV (*i*).

The Simpson index also incorporates richness and evenness but gives greater weight to the most abundant ASVs. Using the definition implemented by `vegan::diversity(index = "simpson")`, it was calculated as

$$1 - \sum_{i=1}^{S} p_i^2.$$

This index represents the probability that two observations selected at random from a sample belong to different ASVs. Values approaching 1 indicate greater diversity, whereas values approaching 0 indicate that the sample is dominated by one or a small number of ASVs.

Observed ASV richness was calculated for each sample as the total number of unique ASVs with a non-zero abundance. This measure describes how many distinct ASVs were detected but does not consider their relative abundances. Because ASVs do not necessarily correspond directly to formally defined microbial species, this metric is referred to here as “observed ASV richness” rather than “species richness.”

Community evenness was quantified using Pielou’s evenness, also known as the Shannon equitability index:

$$J' = \frac{H'}{\ln(S)},$$

where ($H'$) is the Shannon diversity index and ($S$) is the observed ASV richness. The index describes how evenly sequencing reads were distributed among the ASVs detected in a sample. Values approaching 1 indicate that the ASVs occurred at similar relative abundances, whereas lower values indicate that the community was dominated by a smaller number of ASVs. Evenness was not defined for samples containing fewer than two ASVs because ($ln(S)$) is zero when only one ASV is present.

Phylogenetic α-diversity was evaluated using Faith’s Phylogenetic Diversity (Faith’s PD) and Phylogenetic Species Variability (PSV). Unlike Shannon, Simpson, and observed ASV richness, these measures incorporate the evolutionary relationships among the detected ASVs. To calculate them, a phylogenetic tree representing the relationships among all ASV sequences was constructed.

First, the nucleotide sequence of every ASV was extracted and exported in FASTA format using Biostrings v2.74.1. A FASTA file stores each ASV as a named nucleotide sequence. The sequences were then aligned using MUSCLE v5.1 (6). Multiple sequence alignment arranges homologous nucleotide positions into the same columns, allowing sequence differences to be interpreted in an evolutionary context.

The resulting alignment was refined using Gblocks v0.91b (7). Gblocks removes poorly aligned or highly ambiguous regions while retaining nucleotide blocks that can be aligned reliably across the sequences. Removing uncertain alignment positions reduces the likelihood that alignment errors will influence reconstruction of the phylogenetic tree.

A phylogenetic tree was subsequently inferred from the refined alignment using FastTree v2.2 (8). The tips of this tree represent individual ASVs, while its branches represent their inferred evolutionary relationships and sequence divergence. The tree was midpoint-rooted using phangorn v2.12.1 (9). Midpoint rooting places the root halfway along the path between the two most distant tips in the tree. This provides an estimated root when no suitable external reference sequence, or outgroup, is available.

Faith’s PD and PSV were then calculated using Picante v1.8.2 (10). Faith’s PD is the sum of the branch lengths required to connect all ASVs detected in a sample. A higher value therefore indicates that the sample contains ASVs spanning a greater amount of evolutionary history. Because adding ASVs can add branches to the tree, Faith’s PD is generally influenced by observed richness.

PSV measures how phylogenetically related the ASVs within a sample are while reducing the direct influence of richness. Values approaching 1 indicate that the detected ASVs are relatively distantly related, whereas lower values indicate that they are more closely related or phylogenetically clustered. Faith's PD and PSV consequently describe different aspects of phylogenetic diversity: Faith's PD quantifies the total amount of evolutionary history represented, while PSV describes how that evolutionary history is distributed among the detected ASVs.

Differences in α-diversity among the Bangkok, Tak, and Phatthalung populations were evaluated using pairwise Wilcoxon rank-sum tests implemented in rstatix v0.7.2 (11). This non-parametric test compares the distributions of diversity values between two independent groups without assuming that the data are normally distributed. Separate comparisons were performed for Bangkok versus Tak, Bangkok versus Phatthalung, and Tak versus Phatthalung.

Multiple sequence alignment, alignment refinement, and phylogenetic-tree construction were performed from the Bash command line. Sequence extraction, diversity calculations, tree rooting, phylogenetic-diversity calculations, and statistical comparisons were conducted in R v4.4.1 (12).

#### β-diversity

Between-sample microbial diversity (β-diversity) was assessed to determine how strongly microbial community composition differed among samples. Four complementary measures were used: Bray–Curtis dissimilarity, Aitchison distance, unweighted UniFrac distance, and weighted UniFrac distance. These measures emphasize different features of the microbial communities and therefore provide complementary perspectives on compositional variation.

Bray–Curtis dissimilarity was calculated from the ASV relative-abundance matrix using vegan v2.6.8 (5). Bray–Curtis dissimilarity compares samples on the basis of differences in ASV abundances. It ranges from 0 to 1, where 0 indicates that two samples have identical relative-abundance profiles and 1 indicates that they share no ASVs with non-zero abundance. This measure considers the abundance of each ASV but does not incorporate evolutionary relationships among ASVs.

Aitchison distance was used as a complementary abundance-based measure that accounts for the compositional nature of microbiome data. Sequence abundances are compositional because the values within each sample represent proportions of a fixed total; consequently, an increase in the relative abundance of one ASV necessarily affects the relative abundances of the others. To address this dependency, the relative-abundance matrix was transformed using the centered log-ratio (CLR) transformation:

$$\mathrm{clr}(x_i) = \ln\left(\frac{x_i + y}{g(x + y)}\right),$$

where (*x_i*) is the abundance of ASV (*i*) and (*g(x)*) is the geometric mean abundance of all ASVs in the sample. Since division through zero is not defined, we added a pseudocount (*y*) of 0.5 to the counts. Aitchison distances were then calculated as the Euclidean distances between samples in CLR-transformed space using compositions v2.0.8 (13). Thus, the Aitchison distance describes differences in the log-ratios among ASV abundances rather than differences in their raw or relative abundances.

Phylogenetic β-diversity was evaluated using unweighted and weighted UniFrac distances, calculated with phyloseq v1.50.0 (14). The midpoint-rooted phylogenetic tree used for the α-diversity analyses was also used for these calculations. UniFrac measures the proportion of phylogenetic branch length that is unique to either of two microbial communities. Unweighted UniFrac uses only the presence or absence of ASVs and is therefore particularly sensitive to differences involving rare or low-abundance lineages. Weighted UniFrac additionally incorporates ASV abundances, giving greater influence to phylogenetic lineages that are abundant within the communities. The two UniFrac measures therefore distinguish changes in community membership from changes in the abundance of phylogenetic lineages.

The relationships among samples were visualized using ordination methods, which reduce complex, multidimensional community data to a small number of axes. Principal coordinates analysis (PCoA) was performed on the Bray–Curtis, unweighted UniFrac, and weighted UniFrac distance matrices using ape v5.8 (15). PCoA positions samples so that the distances between points approximate their ecological dissimilarities. Samples plotted close together have more similar microbial communities, whereas samples farther apart have more dissimilar communities. The percentage of variation represented by each coordinate was obtained from its corresponding eigenvalue.

Because Aitchison distance is Euclidean distance in CLR-transformed space, principal component analysis (PCA) was performed directly on the CLR-transformed abundance matrix using stats v4.4.1. PCA identifies the major directions of variation in the transformed data. As with PCoA, samples positioned close together in the ordination have more similar community compositions. The proportion of variance explained by each principal component was used to describe how much of the overall compositional variation was represented by each axis.

Differences in within-population compositional variability were assessed using multivariate dispersion analysis. For each β-diversity matrix, the distance from every sample to the centroid of its population—Bangkok, Tak, or Phatthalung—was calculated using vegan. These distances quantify the degree of variation among microbial communities within each population. Larger distances indicate greater within-population heterogeneity, whereas smaller distances indicate that samples from the population have more similar microbial compositions. Differences in mean dispersion among the three populations were evaluated using analysis of variance (ANOVA).

Differences in overall microbial community composition among populations were tested using pairwise permutational multivariate analysis of variance (PERMANOVA), implemented in pairwiseAdonis v0.4.1. PERMANOVA determines whether samples belonging to different populations are more compositionally dissimilar than expected under random reassignment of population labels. Unlike conventional ANOVA, it operates directly on a distance matrix and does not require the original multivariate data to follow a normal distribution.

Separate PERMANOVA comparisons were conducted for Bangkok versus Tak, Bangkok versus Phatthalung, and Tak versus Phatthalung using each of the four β-diversity measures. Statistical significance was assessed using 10,000 permutations. During each permutation, population labels were rearranged and the test statistic was recalculated, producing a null distribution representing the differences expected if population membership were unrelated to microbial composition. P-values from the pairwise comparisons were adjusted using the Bonferroni method (`p.adjust.m = "bonferroni"`) to control the family-wise error rate associated with performing multiple tests.

PERMANOVA and multivariate dispersion address different questions. PERMANOVA evaluates whether the average community composition, or multivariate centroid, differs among populations, whereas dispersion analysis evaluates whether the amount of within-population variation differs. The dispersion results were therefore considered when interpreting PERMANOVA because significant differences in dispersion can contribute to an apparent PERMANOVA difference, particularly when group sizes are unequal. All β-diversity calculations, ordinations, and statistical analyses were conducted in R v4.4.1.

#### Differential abundance analysis

Differential abundance analysis was performed to identify individual microbial taxa whose abundances differed among the Bangkok, Tak, and Phatthalung populations. Analyses were conducted using Analysis of Compositions of Microbiomes with Bias Correction 2 (ANCOM-BC2) in pairwise-comparison mode (16). ANCOM-BC2 is designed for microbiome count data, which are compositional because the number of reads assigned to one taxon depends partly on the total number of reads obtained from the sample. The method estimates and corrects for sampling-fraction bias before testing whether the abundance of each taxon is associated with the variable of interest.

Differential abundance was evaluated separately at the ASV and genus levels. At the ASV level, each exact sequence variant was treated as an individual feature. For the genus-level analysis, ASVs assigned to the same genus were aggregated by summing their read counts within each sample. Genus-level aggregation provides a broader taxonomic summary and may increase the number of observations available for detecting changes shared by related ASVs. It can, however, conceal differences among ASVs belonging to the same genus. This procedure represents taxonomic aggregation rather than sequence clustering, because the ASVs themselves were not reclustered on the basis of sequence similarity.

Before analysis, samples with fewer than 1,000 total reads were removed because their microbial composition was considered insufficiently characterized. Taxa detected in fewer than 10% of the retained samples were also excluded. This prevalence filter removed very rare taxa for which abundance differences could not be estimated reliably and reduced the number of statistical tests performed. The remaining untransformed count data were supplied to ANCOM-BC2.

Population was included as the primary explanatory variable in the statistical model. Sex, age, and body mass index (BMI) were included as additional fixed effects, corresponding to a model of the general form: microbial abundance ~ population + sex + age + BMI.

Including these variables did not remove them from the data. Instead, it allowed differences among populations to be estimated while statistically adjusting for variation associated with sex, age, and BMI. The estimated population effects therefore represent differences in microbial abundance conditional on the other variables included in the model. This adjustment was intended to reduce potential confounding, although it cannot eliminate confounding caused by unmeasured variables.

Pairwise contrasts were used to compare microbial abundances between Phatthalung and Bangkok, Tak and Bangkok, and Tak and Phatthalung. For each taxon and contrast, ANCOM-BC2 estimated the direction and magnitude of the abundance difference as a log-fold change and tested the null hypothesis that no difference was present after adjustment for the model covariates. A positive log-fold change indicated a higher estimated abundance in the first population named in the contrast, whereas a negative value indicated a lower estimated abundance relative to the second population.

Multiple-testing-adjusted P-values were used to determine statistical significance because many taxa were tested simultaneously. Following completion of the full analysis, taxa that were significantly different in at least one of the three pairwise comparisons were retained for visualization and biological interpretation. Filtering for visualization was performed only after statistical testing; all taxa that passed the initial read-depth and prevalence criteria were included in the differential abundance models.

#### Lifestyle analysis

Lifestyle variables with more than 50% zero values or more than 50% missing observations were excluded. Among the remaining variables, features with no variation across samples were removed because they could not contribute to comparisons. Pairwise Pearson correlations were calculated using compositions v2.0.8 (13), and variables strongly associated with locality (($|r|>0.5$)) were excluded to reduce redundancy with the geographical grouping. Non-lifestyle variables—including sex, age, height, BMI, calprotectin, chromogranin, IgM, IgA, genetic admixture, and the inflammatory bowel disease polygenic risk score (`PRS_IBD`)—were also removed manually. The remaining variables were retained as the lifestyle-feature dataset.

Because the lifestyle dataset contained a mixture of continuous, binary, and categorical variables, differences between participants were quantified using Gower distance in cluster v2.1.6. Gower distance scales each variable according to its type and combines the resulting differences into an overall dissimilarity between each pair of participants. The distance matrix was summarized by principal coordinates analysis (PCoA) using ape v5.8 (15).

Associations between individual lifestyle variables and the first two PCoA axes were summarized using Pearson correlation coefficients. Each feature was represented as a vector with coordinates corresponding to its correlations with PCoA1 and PCoA2. Vector length was calculated as ($\sqrt{r_1^2 + r_2^2}$), where ($r_1$) and ($r_2$) are the respective correlations. Longer vectors indicated stronger overall associations with the two-dimensional ordination.

The ability of lifestyle variables to distinguish Bangkok, Tak, and Phatthalung was evaluated using a random-forest classifier implemented in randomForest v4.7.1.2. Missing lifestyle values were imputed using missForest v1.6.1, which predicts missing entries from the other observed variables. Feature-importance scores were extracted from the fitted model to identify the variables that contributed most strongly to locality classification.

Finally, the association between overall lifestyle differences and microbial community composition was assessed using Mantel tests in vegan v2.6.8 (5). These tests correlated the lifestyle Gower-distance matrix with the Bray–Curtis, weighted UniFrac, and unweighted UniFrac distance matrices. Statistical significance was evaluated by permuting the correspondence between participants while preserving the structure of each distance matrix.

#### Host features and microbial association

Associations of lifestyle and other host characteristics with microbial α-diversity were examined separately within the Bangkok, Tak, and Phatthalung populations. Stratification by population was used because relationships between host factors and the microbiome could differ among populations and population identity was highly correlated with lifestyle and host features. Outcomes included observed ASV richness, evenness, Shannon and Simpson diversity, Faith's phylogenetic diversity, and phylogenetic species variability.

For each diversity outcome, predictor selection was performed using least absolute shrinkage and selection operator (LASSOLasso) regression in glmnet v4.1-8 (17–19). LASSO reduces overfitting by shrinking regression coefficients toward zero and excluding predictors with little contribution to the model. Cross-validation was used to select the penalty parameter (λ), which controls the strength of this shrinkage. The selection procedure was repeated 400 times, and host features selected in more than 80% of runs were considered stable predictors.

The retained predictors were subsequently included in an unpenalized multiple linear regression model for each diversity outcome using `lm()` in stats v4.4.1. This provided coefficients on their original, unshrunk scale, together with confidence intervals and conventional significance tests.

The same procedure was used to examine associations between host features and individual ASVs. For these analyses, the CLR-transformed abundance of each ASV was treated as the outcome, and stable predictors were identified through repeated cross-validated LASSOLasso regression followed by unpenalized linear regression.
